# A Framework for Integrating Directed and Undirected Annotations to Build Explanatory Models of cis-eQTL Data

**DOI:** 10.1101/619452

**Authors:** David Lamparter, Rajat Bhatnagar, Katja Hebestreit, T. Grant Belgard, Victor Hanson-Smith

## Abstract

A longstanding goal of regulatory genetics is to understand how variants in genome sequences lead to changes in gene expression. Here we present a method named Bayesian Annotation Guided eQTL Analysis (BAGEA), a variational Bayes framework to model cis-eQTLs using directed and undirected genomic annotations. In a use case, we integrated directed genomic annotations with eQTL summary statistics from tissues of various origins. This analysis revealed epigenetic marks that are relevant for gene expression in different tissues and cell types. We estimated the predictive power of the models that were fitted based on directed genomic annotations. This analysis showed that, depending on the underlying eQTL data used, the directed genomic annotations could predict up to 1.5% of the variance observed in the expression of genes with top nominal eQTL association p-values < 10^−7^. For genes with estimated effect sizes in the top 25% quantile, up to 5% of the expression variance could be predicted. Based on our results, we recommend the use of BAGEA for the analysis of cis-eQTL data to reveal annotations relevant to expression biology.

## 2 Introduction

A longstanding goal in the field of genetics is to accurately predict the phenotypic consequences of any given variant from the genome sequence alone, i.e. to ‘read the genome’[1]. This would help to reveal the phenotypic effects of very rare variants even if their effect is weak. The effects of such variants are typically studied via whole genome sequencing studies. However these studies often have limited statistical power because, by definition, there are few carriers in any sampled population[2].

Recently, progress has been made in predicting epigenetic marks and transcription factor (TF) binding from genome sequence alone; these sequence-based models predict the effect of any given sequence variant on epigenetic marks (and TF binding) [3][4][5][6][7]. The question now is how to extend these models to predict effects on genetically complex phenotypes, such as common diseases. A mechanistic stepping stone between the regulation of epigenetic marks and the regulation of complex phenotypes is the regulation of gene expression, as suggested by the previous observation that disease-causing sequence variants are enriched in gene expression quantitative trait loci (eQTLs)[8][9]. Thus, there is a need for sequence-based models to predict gene expression.

One strategy to build sequence-based models of gene expression is to leverage sequence-based epigenetic mark models. Results of sequence-based models of epigenetic marks can be interpreted as *directed genome annotations*. A genome annotation is defined as a collection of genome regions that have a shared property such as coverage by a particular epigenetic mark, or evolutionary conservation across species. Each region can potentially carry an intensity value. For *directed annotations*, the sign of its intensity value depends on characteristics of the sequence in the region, such as the presence of a specific allele. A simple motivating example is that of a SNP in a TF binding site. In this situation, the TF can have higher binding affinity for one allele versus the other allele. This can cause consistent directional transcriptional effects: the allele inhibiting binding of an activating TF for instance should lead to decreased expression of the target gene. A simple strategy to express this effect as a directional annotation would be to use TF position weight matrices that calculate TF affinity for a given sequence; current and more sophisticated methods express the same relationship using deep neural networks.[3][4][5][6][7].

Methods to evaluate the effect of directed genome annotations on gene expression have recently been proposed[7][10]. Specifically, *Zhou et al.* predicted variant impact without exploiting eQTL data using models that predict expression from chromatin patterns directly[7]. *Reshef et al.* presented a fast method to determine which directed annotations are enriched in variants causal for a given phenotype. However, the method from *Reshef et al.* is geared towards screening and hypothesis testing rather than towards detailed predictive modeling. For instance, the Reshef model does not account for interactions between the effect of an annotation and the distance to the transcription start site (*TSS*).

Here we present a new predictive model of gene expression, named Bayesian Annotation Guided eQTL Analysis (*BAGEA*). *BAGEA* is a variational Bayes modeling framework to analyze eQTLs using both directed and undirected annotations. *BAGEA* can model interactions between these annotations by weighting the impact of the directed annotation based on the undirected annotations. Consequently, *BAGEA* can directly model phenomena relevant to genetic architecture, such as the relatively larger impact of SNPs close to the TSS on directed annotations compared to that of distal SNPS, making BAGEA mores useful for predictive modeling. *BAGEA*’s results are interpretable and highlight genome annotations that are particularly predictive for gene expression. Further, *BAGEA* can model multiple causal SNPs per region. Our software implementation of *BAGEA* can be run on summary statistics using external linkage disequilibrium (LD) information as well as on individual level genotype data directly. Optionally, using a low rank approximation of the LD information improves run-time and decreases *BAGEA*’s memory requirements.

We used *BAGEA* to analyze results from a *cis*-eQTL meta-analysis in human monocytes and from *cis*-eQTL summary statistics derived from tissues of various origins[9][11]. As additional input, we gave the method regulatory impact predictions of common variants on epigenetic marks from a recent deep neural network model[7]. We specified these predictions as directed directional annotations in the method. We show that *BAGEA* highlighted biologically sensible annotations as particularly predictive of eQTLs. Further we estimated the predictive power of the directed annotations for various eQTL data sets. Overall, our results suggest that *BAGEA* is a useful framework to build predictive models of gene expression based on directed annotations, find biologically relevant annotations, and benchmark methods that produce such directed annotations.

## 3 Results

### 3.1 Model Overview

*BAGEA* models gene expression as dependent on SNP genotypes in *cis*. In general, SNP effects on gene expression depend on both directed and undirected annotations (Figure 1a). *BAGEA* builds predictors of gene expression and ranks annotations by their impact on gene expression. For every gene *j*, *BAGEA* takes as input a genotype matrix ***X***_*j*_, an expression vector ***y***_*j*_, annotation matrices ***V***_*j*_, ***F***_*j*_ and ***C***_*j*_. ***X***_*j*_ has dimensions (*n* × *m*_*j*_), where *n* is the number of individuals assayed, and *m*_*j*_ is the number of SNPs in *cis* of gene *j*’s *TSS*. The matrices ***V***_*j*_ ***F***_*j*_ and ***C***_*j*_ are of dimensions (*m*_*j*_ × *s*), (*m*_*j*_ × *q*), and (*m*_*j*_ × *t*) respectively, where *s*, *q* and *t* are the number of annotations used. *BAGEA* models gene expression as a linear combination of SNP genotypes:

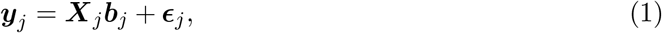

where ***ϵ***_*j*_ is an *i.i.d* normal noise vector and ***b***_*j*_ is a vector of SNP effects. The vector ***b***_*j*_ is modeled as:

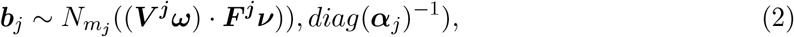

where · is the pointwise product (i.e. Hadamard product) and the *diag*(***x***) operator stands for a diagonal matrix with elements on the diagonal set to ***x***. Detailed descriptions of the terms are as follows:

- ***V***^*j*^ encodes directed annotations. In our applications of *BAGEA*, ***V***^*j*^ is previously computed from sequence-based models, where each column in ***V***^*j*^ represents an epigenetic mark and each row represents a SNP. Each entry in ***V***^*j*^ expresses the predicted effect of a genotype change on the epigenetic mark in question.
- ***F***^*j*^ encodes undirected annotations. Each element in ***F***^*j*^ expresses the presence or absence of the annotation at a SNP’s location. In our applications of *BAGEA*, ***F***^*j*^ is derived from the relative positions of a SNP and gene *j*’s *TSS*, where each column represents a particular region around the *TSS*. For example, if a column in ***F***^*j*^ encodes a region of 20 kilobases (KB) upstream from the *TSS*, all entries for rows corresponding to SNPs within 20 KB upstream of that *TSS* will be set to 1 and entries for all other rows will be set to 0.
- ***ω*** and ***ν*** are vectors that are estimated by *BAGEA*. Specifically, ***ω*** and ***ν*** are the effects of annotations in ***F***^*j*^ and ***V***^*j*^ on the SNP effects ***b***_*j*_.
- 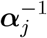 is a vector that is estimated by *BAGEA* and models the variances of elements of ***b***_*j*_. Allowing different variances for the elements of ***b***_*j*_ typically produces sparse estimates where most elements in ***b***_*j*_ are close to zero[12]. Further, ***α*** is modeled as dependent on the undirected annotation matrix ***C***_*j*_. ***C***_*j*_ can potentially be identical to ***F***_*j*_ but can model different undirected annotations as well (see Method Details).

**Figure 1:**
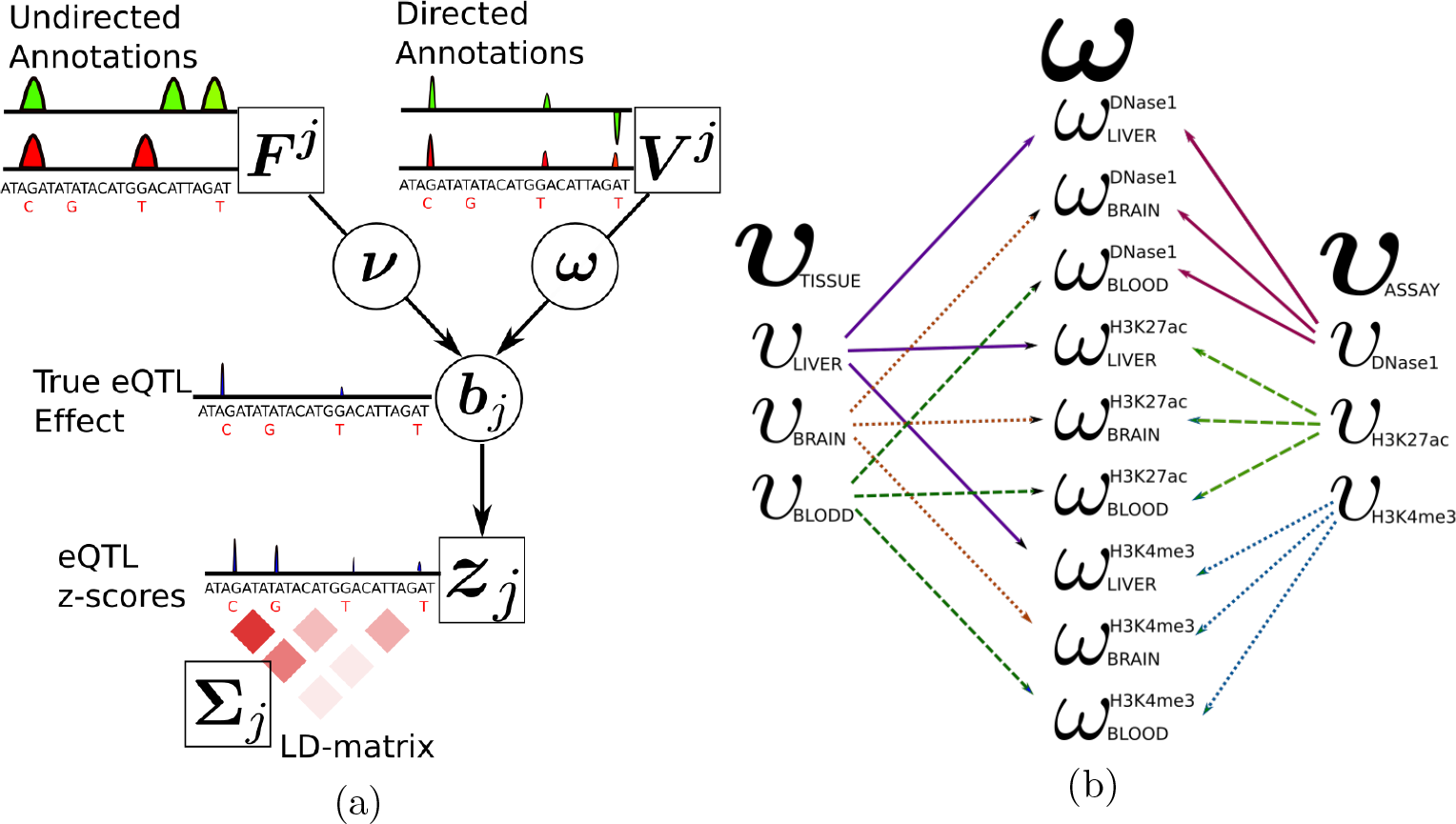
Illustration of BAGEA model components. The core components of the *BAGEA* model in the summary statistics formulation. Observed variables are in squares while estimated variables are circled. Given are ***z***_*j*_, the eQTL z-scores for gene *j*, as well as the LD matrix **Σ***j*, defining the correlation between summary statistics. Further, z-scores are influenced by the true eQTL effects ***b***_*j*_. These effects in turn depend on directed and undirected annotations, ***V***^*j*^ and ***F***^*j*^ respectively. The impact of annotations on ***b***_*j*_ is estimated from the data via ***ω*** and ***ν***. b) An example of the modeling of different priors of elements of ***ω*** using meta-annotations via ***υ*** variable vectors. We assume that directed annotations are available for nine annotations, which were derived from tissues *Liver*, *Blood* and *Brain* via 3 assay types *DNase1*, *H3K27ac* and *H3K4me3*. It is reasonable to assume that for a given eQTL study, particular tissues or cell types are more relevant than others. We model this by introducing a variable *υ* for each tissue (or cell type) that affects the prior distribution of only those elements of ***ω*** that are derived from this tissue, e.g. *υ*_*Liver*_ only affects elements of ***ω*** tied to experiments performed in liver. Analogously to tissue, we model different priors for various for assay types. Shown is the resulting network of influences of the variable ***υ***_*tissue*_, ***υ***_*assay*_ on ***ω***. (For clarity, we used the actual group names as indices, while in the main text, elements of ***υ***’s and ***ω*** are indexed by natural numbers).

Typically, directed annotations are grouped by their cell type or assay type. *BAGEA* can use this grouping structure in order to select groups of annotations that are useful for predicting gene expression (Figure 1b). *BAGEA* selects annotation groups via a modeling strategy that yields sparsity on the annotation group level similar to the group lasso[13]. In *BAGEA*, this grouping strategy is implemented by partitioning annotations into multiple meta-annotations (such as different cell types, assay types etc.). When using this partitioning mechanism, *BAGEA* includes an extra random variable vector ***υ*** of the same length as the number of elements in the partition structure (e.g. the number of cell types, or the number of assay types) (See Methods as well as Figure 1b for an illustrative example). The *k*th element of ***υ***, *υ*_*k*_, controls the variance of the effect sizes for annotations that fall into partitioning group *k*. Specifically, *υ*_*k*_ is proportional to the inverse of the variance of the respective elements of ***ω***. 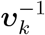 is therefore called the *variance modifier* of annotation partition element *k* (see Methods).

Importantly, the model can be reformulated in terms of the summary statistics 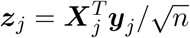 and LD matrices 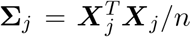. The reformulation enables the application of *BAGEA* to studies for which only summary statistics are available, by estimating **Σ***j* from external sources (see Methods).

### 3.2 Evaluation Strategy for Model Fit

We developed an approach to evaluate the performance of *BAGEA* when fitting directed annotations to genotype and gene expression data. An important feature of *BAGEA* is that its results can be used to predict gene expression for a gene without using any expression data for that gene, but rather using genotypes and genome annotations whose weights are fitted from other genes. We can therefore validate *BAGEA* by training it on gene expression data for one set of genes, and then calculating the extent to which the trained model predicts gene expression for other genes.

We propose a so-called directed predictor 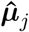, which predicts gene expression for gene *j* based on knowledge of directed annotations and genotype for gene *j*. Using the same notation as in equations (1) and (2), the predictor 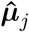 is computed by

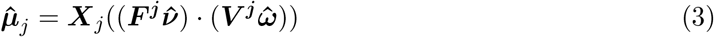

The squared magnitude 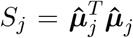 measures how much gene expression variance the model attempts to explain via the predictor 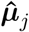. To evaluate the predictor’s accuracy and degree of overfitting, we use the *directed mean squared error* 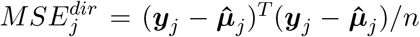. The evaluation of the predictor is performed on a set of genes independent of the ones used to estimate ***ω*** and ***ν***.

Inspection of equation (3) shows that we can reformulate the right hand side in terms summary statistics 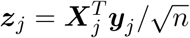, LD matrices 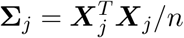, and estimated directed effect of SNPs 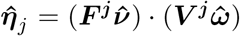, i.e. we can write 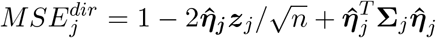 if we assume that ***y***^*T*^***y*** = *n*. In principle, the reformulation allows us to calculate a predictor’s directed mean squared error, even if only summary statistics are available, by approximating **Σ**_*j*_ from external sources.

### 3.3 Directed Annotations Derived from Blood can Partially Explain *cis*-eQTLs in Monocytes

We used *BAGEA* to determine the extent to which annotations can predict gene expression in *CD14* positive monocytes. To this end, we aggregated data from two eQTL studies on expression genetics in *CD14* positive monocytes[14][15]. For directed annotations, we used predictions of genetic variant effects on epigenetic marks (12 different histone mark assays and DNase1 with 4 different peak calling strategies) in various blood-derived cell types from the pre-trained *Expecto* model. *Expecto* is a deep learning framework that predicts epigenetic marks based on sequence context and performs *in silico* mutagenesis to evaluate the consequences of sequence variants[7]. *Expecto* yielded 2002 directed annotations of which 253 were from blood related celltypes. These are referred to as the *Blood* annotation subset in this paper. We partitioned these directed annotations by cell type and assay type, respectively, and modeled separate prior variance terms for each partition (Figure 1b).

To train *BAGEA*, we used gene expression data from human chromosomes 1 through 15, filtering for genes with a top nominal *cis*-eQTL p-value lower than 10^−10^, i.e. only genes that had a SNP in *cis* showing a signficant assocation with a p-value lower than 10^−10^ were included. To test model fit, we predicted expression for genes on chromosomes 16 through 22 with a top nominal *cis*-eQTL p-value below 10^−10^. Specifically, we used the model fit on the training set to derive the estimates 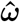 and 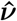(see Equation 2). We then used these estimates to calculate the directed predictors 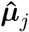 for genes on the test set (see Equation 3). To assess the predictive power of 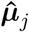, we calculated 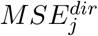 for every gene in the test set. We observed that directed genome annotations can partially explain gene expression variance (Figure 2). The average *MSE*^*dir*^ across all genes was 99.5%, which was significantly smaller than 100% (as evaluated by bootstrap sampling genes; p-value smaller than 10^−4^). 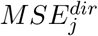 showed a dependence on predictor size *S*_*j*_ (where 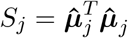), such that for the top quartile of genes when ranked by *S*_*j*_, the directed component was estimated to predict 1% to 3% of expression variance (Figure 2a). For each gene, the variance explained is bounded by the additive genetic variance component in *cis* which is typically much lower than 100%. We estimated the variance of expression explained for each gene in *cis* in an unbiased way via Haseman-Elston (HE) regression[16]. This approach suggested that around 6.6% of the total genetic variance in *cis* was explained by the externally fitted directed component 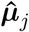 for genes in the top quartile w.r.t *S*_*j*_ (Figure 2b).

**Figure 2:**
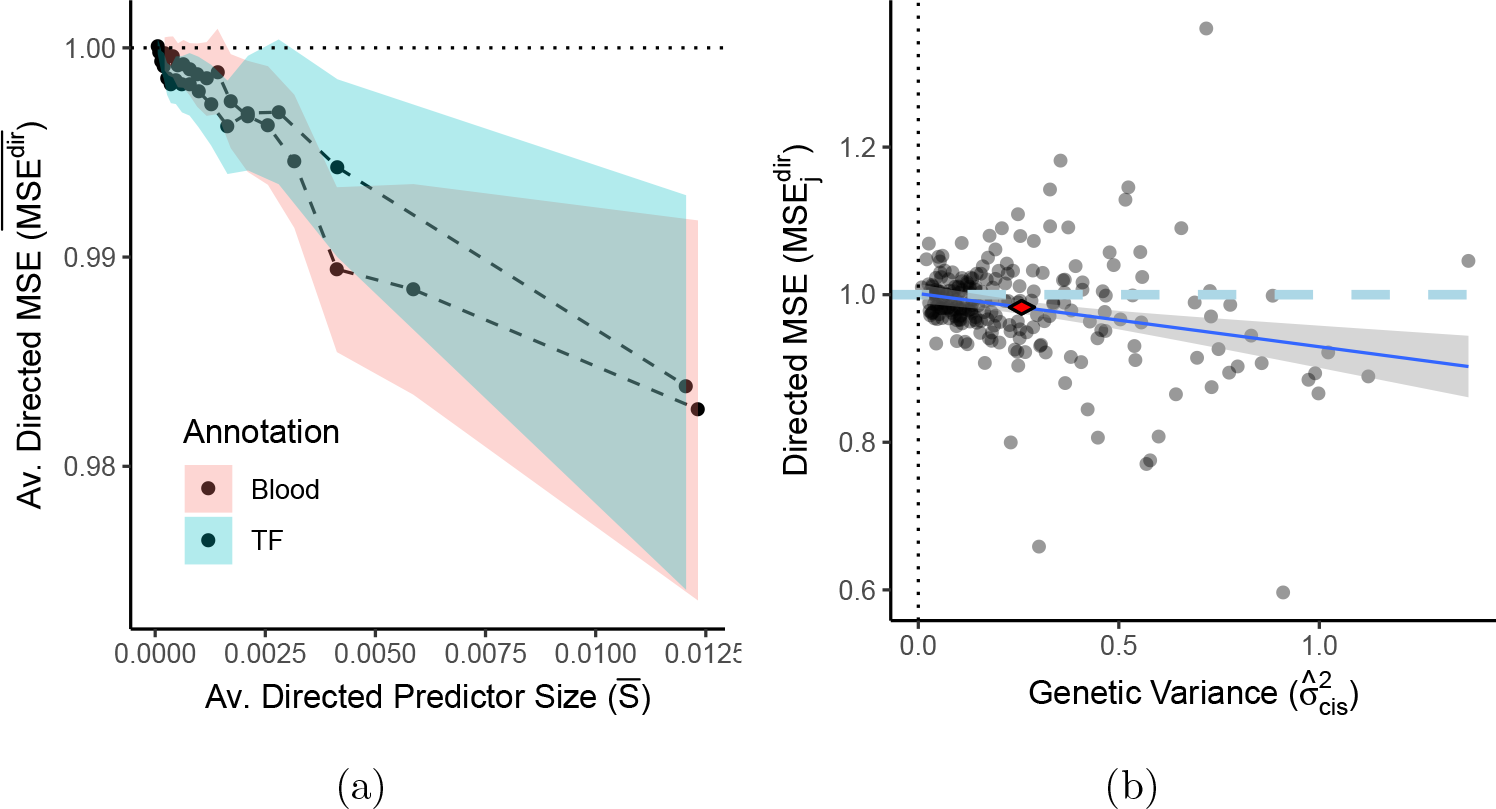
Gene expression variance can be partially explained by directed genome annotations. The BAGEA model was fitted on genes in the training set (all genes on chromosomes 1 through 15) using monocyte eQTL data on genes with a top nominal p-value below 10^−10^, and with Expecto-derived directed annotations. Expecto includes 2002 total annotations, of which one of two subsets were used: 253 annotations derived from histone and DNase1 assays in a blood related cell types (*Blood*), or, alternatively, 690 annotations derived from TF ChIP-Seq (*TF*). For each gene *j* in the test set (all genes on chromosomes 16 through 22 with a top nominal p-value below 10*−*10), we calculated the directed predictor of expression 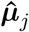. As a measure of a predictor’s size, we use its squared magnitude 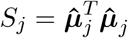. To evaluate the predictor’s performance, we calculated 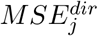, the mean squared error (*MSE*) when predicting gene expression ***y***_*j*_ from 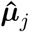. To estimate what the smallest attainable 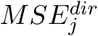 would be, we estimated 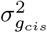, the additive genetic variance in *cis* via Haseman-Elston regression per gene. a) The relationship between the MSE of the predictor and its squared magnitude. We sorted results by predictor Size *S*_*j*_ and averaged 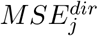 within a sliding window containing 25% of genes and step size of 5% of data. **Averaged Directed Predictor Size** 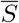: The mean value of *S*_*j*_ per window on the horizontal axis; **Averaged Directed MSE** 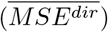: The averaged 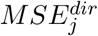 of genes falling into the window on the vertical axis. The 95% confidence interval for each window was derived by bootstrapping. Most variance is explained by genes in the top quartile when ranked by *S*_*j*_. b) The relationship between 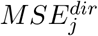 and 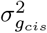 for genes in the top quartile when ranked by *S*_*j*_. **Genetic Variance** 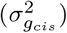: The estimated additive genetic variance in *cis* on the horizontal axis. **Directed MSE** 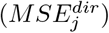 on the vertical axis. 95% confidence intervals for the mean of both the *MSE*^*dir*^ and 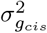 are represented as the corners of the red diamond (i.e. the confidence interval for the average *MSEdir* is given by the upper and lower corner, whereas the confidence interval for the average 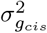 is given by the right and left corner respectively). A linear regression is plotted as the blue line, with 95% confidence interval shown in grey.

### 3.4 Joint Modeling of cis-eQTLs and Directed Annotations Highlights Biologically Relevant Epigenetic Marks

We next evaluated if *BAGEA* can effectively be used to discover which annotations, or groups of annotations, are most predictive of gene expression. We grouped the directed annotations by cell type and assay type, and for each set of annotation groups, we modeled separate prior variance modifiers ***υ***^−1^ (Figure 1b). For each annotation group *k* we measured its contribution to gene expression as its estimated variance modifier 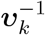 (See Model Overview).

For the monocyte data, *BAGEA* estimated the largest variance modifiers for annotations from *DNase1* as well as *H3K27ac* and *H3K4me3* assays (Figure 3a). This observation is consistent with results from a previous method, using undirected annotations, suggesting that SNPs with an effect on gene expression are enriched in open chromatin (*DNase1*), activated enhancers and promoters (*H3K27Ac*, *H3K4me3*)[17]. Across cell type annotations, *BAGEA* estimated the largest variance modifiers for annotations from two blood cell types that were both *CD14* positive (Figure 3b). This observation matches our expectations because the cells in the underlying expression data were derived from *CD14* positive cells[14][15]. Across all tested pairs of assays and cell types, *BAGEA* estimated the largest positive effect sizes for annotations from *DNase1*, *H3K27ac*,*H3K4me3* assays in *CD14* positive cells (Figure 3c).

**Figure 3:**
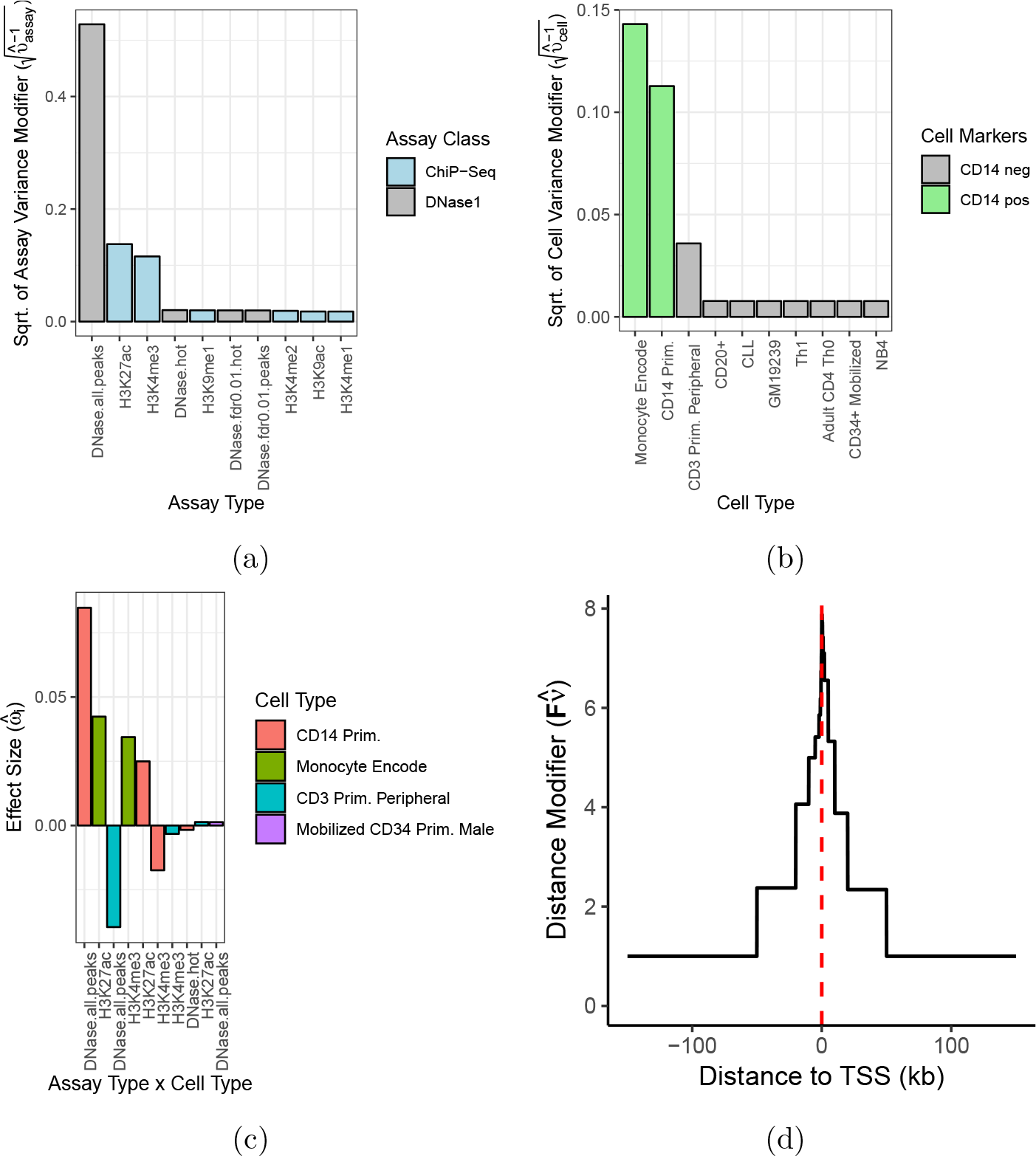
BAGEA, fitted on monocyte eQTL data, selects relevant epigenetic marks and increases directional effect sizes for SNPs close to a TSS. Parameter estimates when applying *BAGEA* to monocyte eQTL data using as directed annotations histone and DNase1 *Expecto* predictions derived from blood-related cell types (i.e. *Blood* from Figure 2). a) For each chromatin assay type, *BAGEA* models an **assay variance modifier** 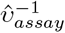 that captures the extent to which that assay type is predictive of gene expression. Shown are the square roots for the assay types with the ten highest variance modifiers (from 17 assay types total). In the *BAGEA* model, *DNase1*, *H3K27Ac* and *H3K4me3* assays have largest modifiers. b) For each cell type, *BAGEA* models a **celltype variance modifier** 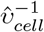, similar to the assay variance modifier in panel a. Shown are the square roots for the cell types with the ten highest variance modifiers (out of 61 cell types). In the *BAGEA* model, *CD14* positive cells have the largest modifiers. c) *BAGEA* reveals which experiments underlying the directed annotations that were most predictive of gene expression. **Assay Type x Cell Type**: Each experiment is a particular assay type performed in a particular cell type. **Effect Size** (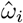, for experiment i): The BAGEA-estimated effect on gene expression. Shown are the ten largest directed annotation effect sizes. In the *BAGEA* model, the experiments using *DNase1*, *H3k27Ac* and *H3Kme4* with *CD14* positive cells have the largest effect sizes. We also see that most of the 253 annotations are estimated to have a close to zero effect. d) Shown is the estimated **distance modifier** of the directed component, 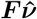. We see a characteristic peak around the *TSS*, implying that the directed annotations are upweighted close to the *TSS*.

It is well known that eQTLs increase in intensity closer to the *TSS*. This suggests that the effects of directed annotations might also be bigger for SNPs close to the *TSS* than for SNPs that are distal. *BAGEA* models SNP distance dependence of directed annotation effects by weighting the directed annotation effect term ***V***^*j*^*ω* across SNP’s, with a distance modifier ***F***^*j*^*ν* (see Model Overview). We next tested whether *BAGEA* estimated directed annotation effect sizes to be dependent on a SNP’s distance to the *TSS*. We examined the value of a SNP’s estimated distance modifier 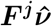 against its position relative to the *TSS*. We observed a characteristic peak around the *TSS* (Figure 3d), suggesting that *BAGEA* can indeed produce a similar pattern of distance dependence for the effect sizes derived from directed annotations as for the eQTL effect sizes themselves.

We repeated this analysis with a different set of directed annotations, namely 690 *Expecto* annotations derived from transcription factor (*TF*) ChIP-Seq in any cell type. We estimated the *TF* annotation subset to be similarly predictive of gene expression as the *Blood* annotation subset (Figure 2a). Parameter estimates for ***ω*** suggest that binding sites of the TF *c-Myc* in cell line *NB4* have the largest effect size on gene expression among all tested 690 annotations (Supplementary Figure 1). *NB4* is a promyelocytic leukemia cell line that can be differentiated into neutrophils or monocytes[18]. *NB4* is therefore expected to have similar expression genetics as *CD14* positive monocytes, and, given that no *TF* ChIP-Seq experiment was performed in monocyte cell lines directly, the large ***ω*** values for *NB4* data are consistent with our expectations.

### 3.5 Modeling Directional Components is Robust to the Use of Summary Statistics

In many cases it is not feasible to compute LD for the population from which the summary statistics were derived (i.e., the study population), and LD has to be derived from other sources (i.e., external genotypes) [19][20]. The use of external genotypes allows publicly available summary statistics to be analyzed without access to restricted individual level genotype data[9]. However, LD computed on external genotypes can only approximate LD patterns of the study population. We therefore need to test the accuracy of methods when using external genotypes.

We evaluated if directed annotation effects were robust to the genetic source of LD information. We used 1000 Genomes data to compute LD as it is publicly-available and widely used for this purpose[21]. We re-fit the *BAGEA* model to monocyte data with the *Blood* an notation subset, using LD matrices derived from European 1000 Genomes data. We then compared ***ω*** estimates when using LD from 1000 Genomes to ***ω*** estimates when using LD from the monocyte data itself, for every annotation in the monocyte Blood data. We observed that the two approaches produced similar effect sizes with a linear regression *R*^2^ of 97.5% and regression slope of 0.96 (Figure 4a). This suggests that directed annotation effect estimates are robust to the source of LD information.

**Figure 4:**
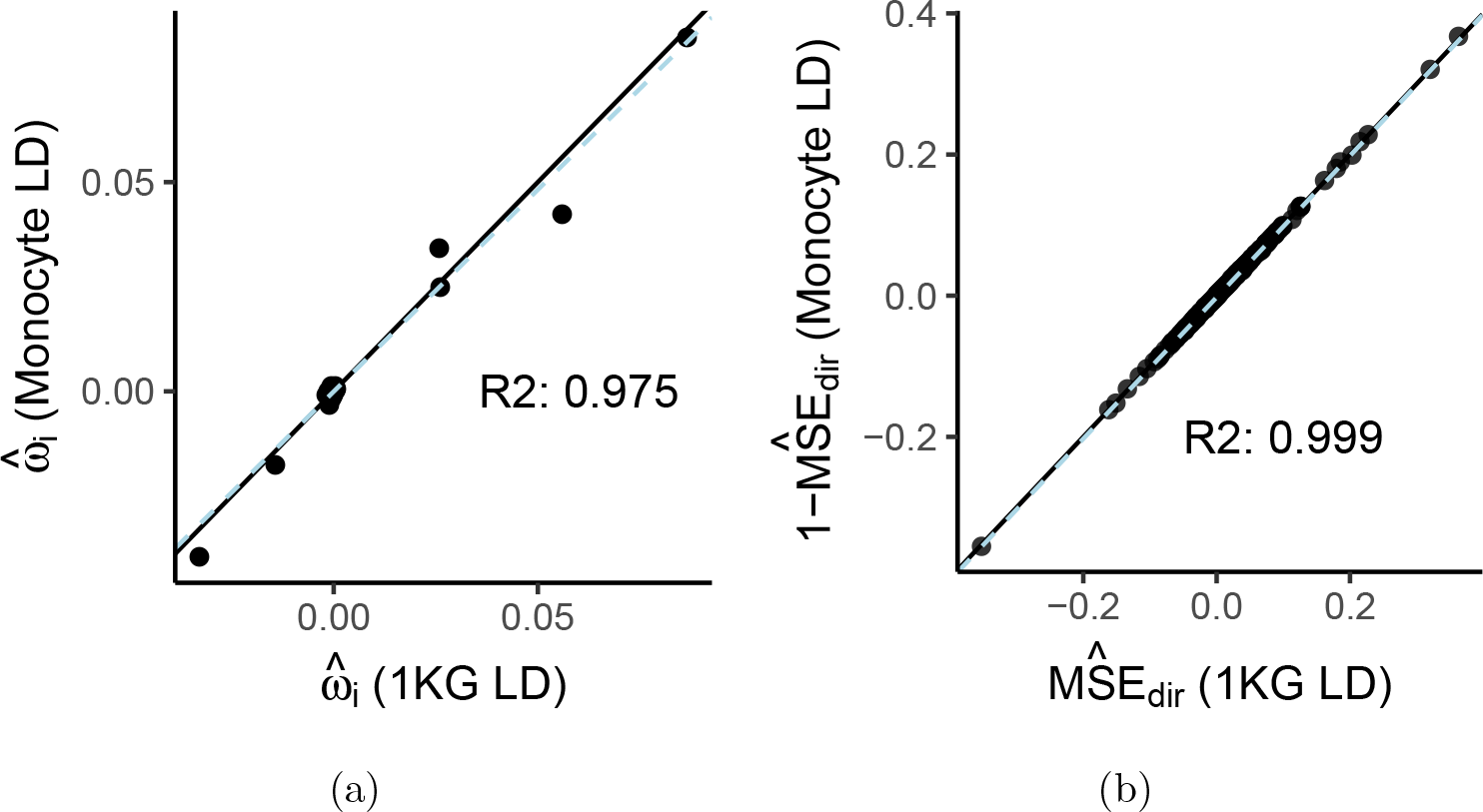
Directed annotation effect estimates and modeling error are robust to source of LD information. a) We sought to investigate the impact on the estimate of the directed annotation effect vector ***ω*** when using cis-eQTL summary statistics with external reference LD information. We retrained BAGEA with the blood monocyte summary statistics using reference LD matrices from the 1000 Genomes Project (1KG). 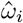 **(1KG)**: Directed annotation effect, measured as ***ω*** estimates from BAGEA using 1KG reference LD information. 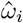 **(Monocyte LD)**: Directed annotation effect, measured as ***ω*** estimates from BAGEA using individual-level genotypes from the monocyte data itself (i.e. using the same genotypes as for the deriving the summary statistics). b) To investigate the extent to which 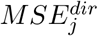 can be approximated using summary statistics and reference 1KG LD matrices, we calculated 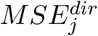 on chromosomes 16 to 22 from summary statistics of monocyte cis-eQTLs (see formula in main text). We then compared these to the original 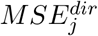 values that were computed using genotypes of the monocyte data sets. The same SNPs were used in both calculations. *R*^2^: The coefficient of determination, measuring goodness-of-fit, from a linear regression of the data shown.

We then explored if the source of LD information affected our estimates of directed mean squared error (*MSE*^*dir*^). To this end, we estimated *MSE*^*dir*^ on chromosomes 16 through 22 from summary statistics and external LD matrices derived from 1000 Genomes alone, and then compared these *MSE*^*dir*^ values to the original *MSE*^*dir*^ values computed with LD derived from monocyte data. We ensured that the same SNPs were included, by removing SNPs with low minor allele frequency (MAF) in either of the sets. We observed that the two sources of LD produced *MSE*^*dir*^ values that agree with each other, with a linear regression R2 of 99.9% and regression slope of 1.002 (Figure 4a).

### 3.6 Analysis of GTEx Summary Statistics Highlights Annotations Gathered from Relevant Tissues

Having established that *BAGEA* performs well when using summary statistics, we next determined if *BAGEA* can identify relevant directed annotations for empirical data for which summary statistics are available but genotypes are not. Specifically, we fit *BAGEA* on summary statistics for eQTL studies of 13 tissues produced by the GTEx consortium with a sample size of at least 300 for each study[9]. We additionally supplemented this set with results for Lymphoblastoid cell lines (LCL) derived from a meta-analysis of GTEx and GEAUVADIS[11]. Because GTEx gathered eQTLs in complex tissues and sampled fewer individuals than were sampled in the monocyte studies, we expected lower power to produce robust parameter estimates. We therefore used different parameter values than in our monocyte analysis, including genes with top nominal *cis*-eQTL p-value lower than 10^−7^. We fitted models using *Expecto* derived annotation for all 1187 histone or DNase1 annotations derived from Roadmap consortium data[22].

We again split genes into training and test set, fitting *BAGEA* on the training set and building directed expression predictors 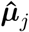 for all genes in the test set. We observed that the average *MSE*^*dir*^ per data set was variable across GTEx data sets ranging from 100% to below 98.5%(Figure 5a). We again saw that increased directed predictor magnitude tended to decrease *MSE*^*dir*^. For instance in fibroblast, the quarter of the genes with the highest directed predictor magnitude had an average *MSE*^*dir*^ of 97.2%, whereas the quarter with the lowest directed predictor magnitude had an average *MSE*^*dir*^ close to 100% (Figure 5b).

**Figure 5:**
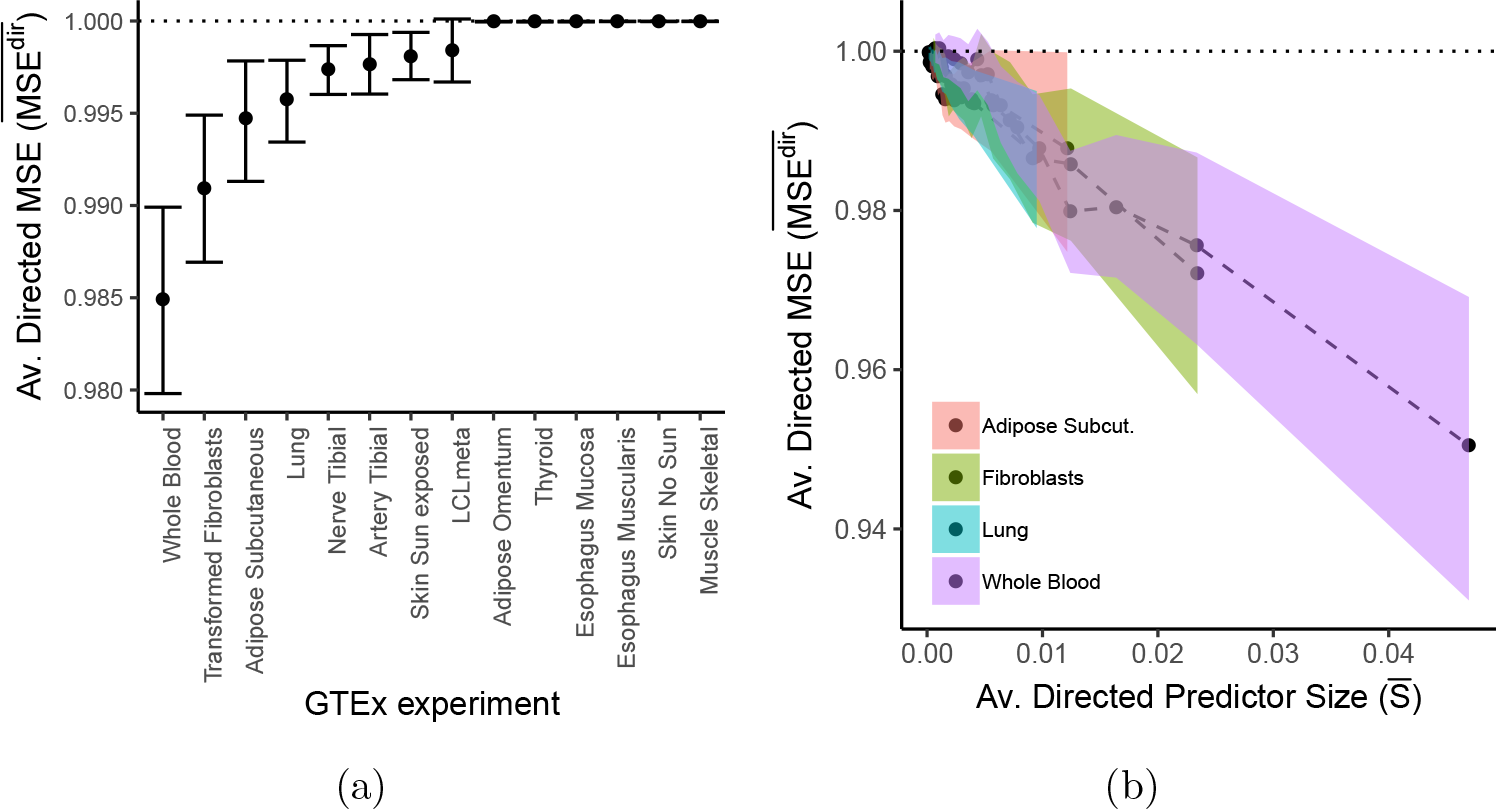
Directed annotations parially explain gene expression variance in GTEx. The BAGEA model was fit using various GTEx eQTL data (supplemented with GEAU-VADIS eQTL data) and with Expecto-derived directed annotations on genes in the trainig set (chr1,..,chr15) with a top nominal p-value< 10^−7^. Expecto includes 2002 total annotations, of which histone and DNase1 annotations from Roadmap were used (1187 annotations in total). For each gene *j* in the test set (chr16,..,chr22 and top nominal p-value< 10^−7^), we calculated an approximate version of *S*_*j*_, the squared magnitude of the directed predictor 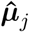, where the approximation uses external LD information. Further, we calculated an approximate version of 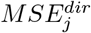, the mean squared error (*MSE*) when predicting gene expression ***y***_*j*_ from 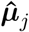. a) Displayed is the average (approximated) 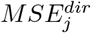 across all genes for each GTEx experiment. 95% Confidence intervals are computed by bootstrapping. b) Displayed is the relationship between the MSE of the predictor and its squared magnitude for the four GTEx experiments with the lowest average 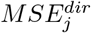. We sorted results by predictor size *S*_*j*_ and averaged 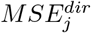 within a sliding window containing 25% of genes within the window and step size of 5% of data. **Averaged Directed Predictor Size** 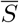: The mean value of *S*_*j*_ per window on the horizontal axis; **Averaged Directed MSE** 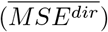: The averaged 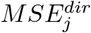 of genes falling into the window on the vertical axis. The 95% confidence interval for each window was derived by bootstrapping. We see that most variance is explained by genes in the top quartile w.r.t. *S*_*j*_.

To mitigate the impact of limited power during variable selection, we additionally fit models without splitting chromosomes into test and validation sets. The distribution of effect sizes of the directional annotations revealed a bias towards positive values (Supplementary Figure 2). Focusing on the largest positive effect sizes(top ten or 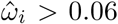), we saw many biologically plausible pairings between the tissue assayed by GTEx via eQTL and the tissue assayed by roadmap for epigenetic marks (Figure 6a). While some of the pairings are obvious from the annotation names themselves (such as correct pairings for lymphoblastoid cells, lung and adipose tissues) others are less obvious yet still plausible. For instance bone marrow derived mesenchymal stem cells (*BMD MSC*) are paired with fibroblast. A recent study found no functional differences between the two cell types leading the authors to support a longstanding opinion in the field that these two cell types should be classified as the same[23][24]. The pairing between Esophagus Mucosa and keratinocytes can be explained by the fact that the Esophagus Mucosa is mainly composed of squamous cells, i.e. keratinocytes[25][26]. The pairing between tibial artery and *BMD MSC* can be explained by the fact that fibroblasts are the main component of vascular adventitia[27]. Our model also paired tibial nerve and muscle, which seems physiologically the least plausible among the ten pairings. When looking at the largest negative values, we saw some of the same tissue pairings repeated, with only one pairing with effect size 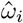 smaller than −0.06 for the pairing between fibroblasts and *BMD MSC*) (Supplementary Figure 3). When looking at the variance modifier estimates for the different assay types, we saw that *DNAse1* and *H3K27ac* epigenetic marks were ranked consistently highly (Figure 6b). Interestingly, among various annotations derived from the same *DNase1* experiments, some performed consistently better than others: *DNase1* peak call annotations outperformed *DNase1* hotspots calls.

**Figure 6:**
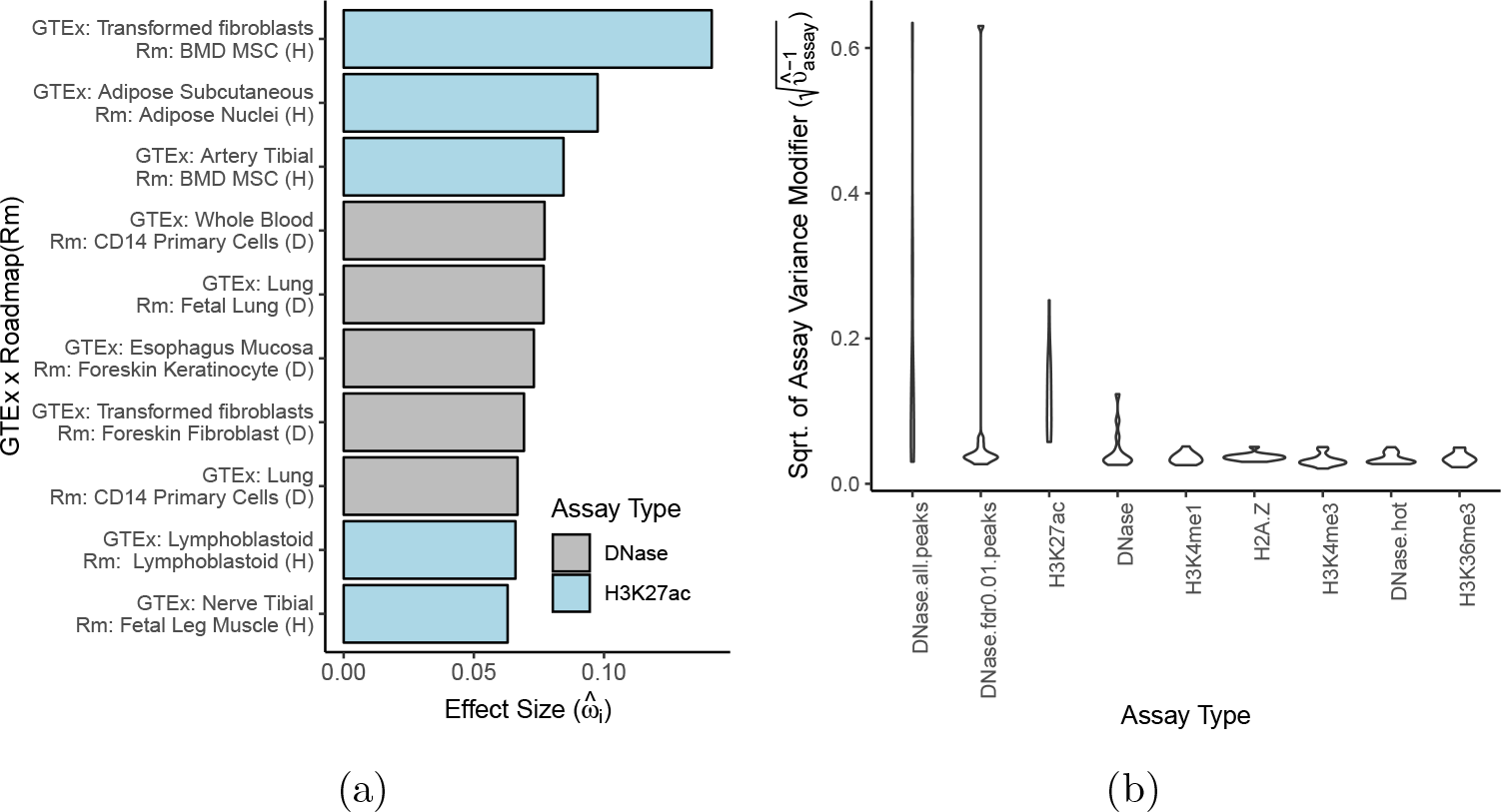
Model fit for GTEx summary statistics highlights directional annotations mainly from plausible cell types. Shown here are various parameter estimates from fitting 13 different GTEx eQTL summary statistics data (supplemented with GEAUVADIS eQTL data) using histone and DNase1 *Expecto* predictions derived from Roadmap (1187 annotations). 4 a) *BAGEA* reveals the experiments underlying the directed annotations that are most predictive of gene expression. **GTEx x Roadmap(Rm)**: Each GTEx eQTL data set highlights particular Roadmap annotations. Shown here are the 10 largest positive effect sizes across all eQTL and annotation pairings. **Effect Size**: The estimate of 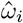 for experiment *i*. 4 b) For each chromatin assay type, BAGEA models an assay variance modifier 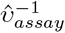 that expresses the extent to which that assay type is predictive of gene expression. Shown here is the distribution of the square roots of the assay variance modifier for any given assay type across all 13 GTEx eQTL data sets. Results are sorted by the maximal value achieved for each assay type and only the 10 highest scoring assay types are shown.

## 4 Discussion

Here we introduced a new method, named *Bayesian Annotation Guided eQTL Analysis* (*BAGEA*). *BAGEA* integrates directed and undirected genome annotations with eQTL data in a variational Bayesian framework to build predictive models of gene expression. We applied this method to eQTL results from *CD14* positive monocytes as follows: First, we derived directed annotations by predicting functional impacts on epigenetic marks for all common SNPs using the pre-trained *Expecto* deep neural net[7]. Second, from these *Expecto* results, we extracted two annotation subsets of particular interest: histone ChIP-Seq and DNase1 in blood-derived cell types (the *Blood* annotation subset), and TF ChIP-Seq in any cell type (the *TF* annotation subset). We then ran *BAGEA* on both annotation subsets separately, while allowing the effect of the directed annotations to depend on the distance to the *TSS*. We tested whether the model had explanatory power with a training and test protocol (i.e. explanatory power was estimated on genes that were excluded from training). We saw that the directed component ***μ*** of the model explained part of the gene expression variance in a statistically significant manner (Figure 2a). For genes with a strong cis-eQTL (p-value< 10^−10^) and in the top quartile for ***μ***^***T***^ ***μ***, we estimated that the *Blood* derived directed component explained 6.6% of total additive genetic variance in *cis* (Figure 2b). Importantly, *BAGEA* prioritized annotations that cohere with widely accepted biological knowledge and are supported by existing literature (Figure 3).

Next, we investigated to which extent the model fit was affected when the LD information was approximated via reference genomes. We observed agreement between the results in terms of the directed component, suggesting that the use of eQTL summary statistics together with external LD data is justified (Figure 4). We therefore used *BAGEA* to analyze eQTL summary statistics results from GTEx. To accommodate the wide range of tissues explored in GTEx, we expanded the number of directed annotations used in the fitting process to over a thousand. While for some tissues, the analysis strategy was underpowered to derive a predictive model of gene expression from directed annotations, others had a significant fraction of gene expression explained by directed annotations (Figure 5). Many of the directed annotations *BAGEA* selected, were derived from tissues that were biologically related to the original tissue of the eQTL studies (Figure 6a). Additionally, we observed that *DNAse1* and *H3K27ac* epigenetic marks were selected across many different eQTL studies (Figure 6b).

BAGEA belongs to a class of models that allow the prior probability distribution of a SNP’s effect size to vary based on the genome annotations with which it overlaps[17][28][29]. These prior models explored the impact of undirected annotations. While *BAGEA* can model undirected annotations, the main novelty comes from the concomitant modeling of directed and undirected annotations as well as interactions thereof. Using directed annotations to explain natural variation in phenotypes was also recently proposed by both *Zou et al.* and *Reshef et al.*, albeit with different modeling philosophies[7][10]. *Zou et al.* use a model that predicts expression from chromatin patterns directly. This has the advantage that genotype data is not needed. However, this method does not model the causal impact of epigenetic marks on expression levels but rather correlations, potentially negatively impacting modeling accuracy. *Reshef et al.*’s LD profile regression method has more similarities to *BAGEA* as it can also be used to analyze directed annotations and eQTL summary statistics. However, the method is geared towards multiple hypothesis testing rather than high predictive accuracy. Compared to *BAGEA*, the fitted model is simpler allowing for fast analysis of large collections of data. The increased speed comes at the cost of not being able to model certain features like interactions of directed and undirected annotations (such as distance to *TSS*). *BAGEA* uses a modeling approach that has both prediction and interpretability in mind. It allows for more complex model features while it is still useful for revealing relevant biology. Indeed, when using BAGEA on various eQTL datasets, *BAGEA* highlighted many relevant cell types. Further, allowing the directed component to depend on the distance to the *TSS* improved the model fit.

There are at least two drawbacks to *BAGEA*’s model complexity. First, there is a substantial computational cost to fit the model. To mitigate this issue, we used computational tricks such as fast matrix inversion of approximated LD matrices and parallelization. Second, variational model fitting approach does not provide confidence intervals. While it does provide credibility intervals, the approximative nature of mean field variational inference makes these credibility intervals often unreliable[30]. In our analysis, we opted for evaluating statistical significance of the model results by using a training and test protocol.

Future research could investigate whether using a different variational approximation rather than the mean field approximation provides better estimates of the true credibility intervals. We estimated the extent to which epigenetic marks are able to predict the genetic component of gene expression in *cis*. Our results show that while the current generation of directed annotations can partially explain the genetic *cis* component of gene expression, most of the genetic *cis* component remains unexplained, indicating that there is still room for improvement. Future gains in this space will likely come from both improved directed annotations as well as improved modeling.

## 5 Methods

### 5.0.1 Model Details

We assume individual level genotype and expression data for *n* individuals. For gene *j*, we model its *n* × 1 expression vector ***y***_*j*_ as

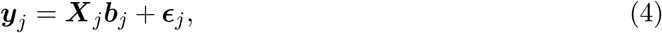

where ***X***_*j*_ is the *n* × *m*_*j*_ genotype matrix for the *m*_*j*_ SNPs surrounding gene *j*’s *TSS*. ***b***_*j*_ is the *m*_*j*_ × 1 vector of SNP effect sizes and ***ϵ***_*j*_ the expression noise unexplained by the genotype.

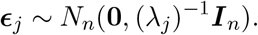

The noise term precision *λ*_*j*_ is modeled in a hierarchical fashion:

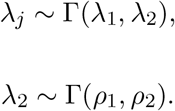

with hyperparameters *λ*_1_, *ρ*_1_ and *ρ*_2_ (while this notation is overloaded, we expect it is clear from context which parameter is meant). We model the vector of effect sizes ***b***_*j*_ as a multivariate normal, whose mean and covariance is affected by annotation matrices. For gene *j* we assume undirected 0 − 1 coded annotation matrix ***F***^*j*^ and a directed continuous annotation matrix ***V***^*j*^, with dimensions *m*_*j*_ × *q* and *m*_*j*_ × *s* respectively. Then,

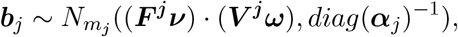

with ***α***_*j*_ being a vector of independently drawn gamma distributed random variables (the modeling is described further down). ***ω*** and ***ν*** are *s* and *q* dimensional multivariate normal distributed random variables respectively. ***ω*** denotes the vector of activities of directed annotations, whereas ***ν*** allows the overall weight that the directed annotations contribute to the effect size vary based on undirected annotations. This allows, for instance, the impact of the directed annotations to vary dependent on the distance to the *TSS*. ***ω*** is modeled in a hierarchical fashion

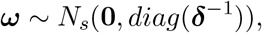

where ***δ*** is again modeled as a random variable. The choice of model for ***δ*** enables the implementation of a grouping structure on the directional annotations (in our application, these groupings are the assay used to derive the annotation and the cell type in which the assay was performed). We allow the model to fit differences in prior variances based on group membership. Thereby, entire groups of directional annotation effects are shrunk to zero (akin to the group lasso[13]). Let ***d***^*j*^ be a positive integer vector of length *s* taking *h*_*j*_ different values, i.e ***d***^*j*^ partitions the vector of directed annotations into *h*_*j*_ groups (in this context, *j* = 1,.., *w* runs over the meta-annotations, e.g. if the modeled meta-annotations are cell type and assay type, *j* can either take the value one or two). Let ***υ***^*j*^ be a random vector of length *h*_*j*_ (i.e. these are the group specific weights). Then,

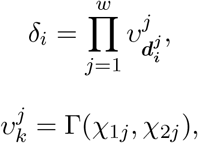

with hyperparameter *χ*_1*j*_. *χ*_2*j*_ is modeled as

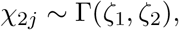

with hyperparameters *ζ*_1_ and *ζ*_2_.

***ν*** is modeled as

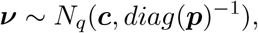

where ***p*** and ***c*** are hyperparameter vectors of length *q*.

The vector of precisions of the effect size vector ***α*** is modeled as

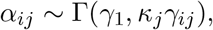

where *γ*_1_ is a hyperparameter. Note that letting the precision for each SNP vary leads to sparse estimates for ***b***_*j*_; this is akin to automatic relevance determination (ARD) regression[12]. *κ*_*j*_ is a genewise parameter modeled in a hierarchical fashion

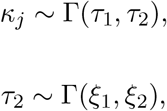

where *τ*_1_, *ξ*_1_ and *ξ*_2_ are hyperparameters. To model *γ*_*ij*_, we again make use of annotation matrices. For gene *j* assume undirected 0 − 1 coded annotation matrix ***C***^*j*^ of dimension *m* × *t*. Then the SNP-wise precision modifier *γ*_*ij*_ is modeled as

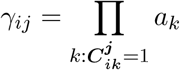

where 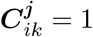 if annotation *k* is active at index *i* in gene region *j*. Further,

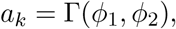

where *φ*_1_ and *φ*_2_ are hyperparameters.

### 5.1 summary statistics adaptation

Instead of using individual level genotype and expression data, we can reformulate the model for the use of summary statistics. Multiplying equation 4 with 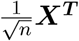 gives

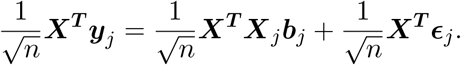

A natural model to use with summary statistics is therefore,

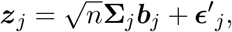

where ***z***_*j*_ is the vector of summary statistics, **Σ**_*j*_ is the LD matrix and 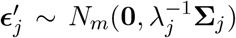 **Σ**_*j*_ can be approximated from external sources such as 1KG[21]. Alternatively, we can use an approximate and regularized version of the empirical LD matrix (see below).

### 5.2 Model fitting

The model was fit using variational bayes approach[30]. As the model is in the conjugate exponential family, we can use the variational message passing strategy[31]. For detailed updating steps see the supplementary information. Naive updates can be prohibitively expensive due to the requirement to invert many large matrices of the form (*c****X***^***T***^***X*** + ***D_α_***), where *c* is a constant and ***D_α_*** is a diagonal matrix. To speed up computation, we can approximate the LD matrix *c****X***^***T***^***X*** with a low rank approximation 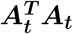, where ***A***_***t***_ is a *t* × *m* matrix with *t < m*. This allows us to speed up a time critical matrix inversion step.

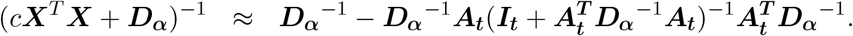

If ***X*** is already low rank, it is computationally advantageous to use an 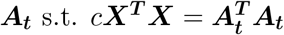. If 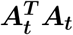 deviates from *c****X***^***T***^***X***, we need to use the summary statistics formulation to avoid convergence issues. For more detail, see the supplementary information.

### 5.3 Deriving Annotations

For common SNPs (minor allele frequency (MAF) above 2.5% in the 1000 Genomes European population[21]), we ran the *Expecto* model to predict the effect of the variant on epigenetic marks[7]. For each SNP we predicted the epigenetic effects within the 200 bp region encompassing it. For most SNPs the effects are very close to zero, allowing us to sparsify the results. Absolute effects smaller than 0.008 were set to zero and all other effects were shrunk towards zero by 0.008 via *x*_*new*_ = *x* − 0.008 · *sgn*(*x*). Next, results for both strands were averaged and the shrinking procedure repeated with a threshold of 0.008. This yielded matrix with 98.4% of entries zero. The directed annotations were then scaled to have all the same 2-norm. The magnitude of the 2-norm was set to the average of the unscaled 2-norms. These were the directed annotations used in *BAGEA*.

For undirected annotations, we used upstream and downstream distances to the *TSS*. Distance to *TSS* annotations as well as SNP positional annotations were downloaded from the UCSC genome annotation database with SNP and gene annotations taken from the *refGene* and *snp147Common* tables respectively (see link below)[32].

### 5.4 cis-eQTL data sets

For monocyte eQTL data, we used two preprocessed monocyte datasets with a combined sample size of 1176 (418 from *Fairfax et al.* and 758 from *Rotival et al.* respectively)[14][15]. Expression matrices were quantile normalized and 10 PEER factors as well as 5 genotype PCs removed[33]. Genotype data was quality control filtered (4% SNP level missingness; 5 % individual level missingness; Hardy-Weinberg p-value above 10^−13^ relatedness below 0.1875) and imputed using the human genome reference panel[34].

We further downloaded eQTL summary statistics for various tissues produced by the *GTEx* project if the number of samples was above 300 individuals[9]. Additionally, for LCL, we meta-analyzed eQTL summary statistics released for 117 samples by *GTEx* with summary statistics derived from 358 European PEER-controlled samples collected as part of the *GEUVADIS* study[11].

### 5.5 Running *BAGEA*

For the monocyte eQTL analysis, *BAGEA* was run with default hyperparameter settings (see supplementary information). Genotypes within a window of 150KB around a gene’s *TSS* were used to construct a genewise LD matrix. Each Genewise LD matrix was approximated via singular value decompostion with a low rank symmetric matrix of equal top eigenvalues and eigenvectors, such that the trace of the approximation matrix was at least 99% of the trace of the original LD matrix. Then, a scaled identity matrix was added such that the trace of the resulting matrix was equal to the trace of the original LD matrix. As undirected annotations, distance windows around the TSS (50KB, 20KB, 10KB, 5KB, 2KB, 1KB, 0.5KB, 0.25KB) split into upstream and downstream windows were used. For all GTEx summary statistics analysis, reference 1KG LD matrices where calculated and replaced with low rank approximation with 95% of the matrix trace kept, anlagously to the above procedure. Default hyperparameter settings where used except for *c* which was set to *0.3* instead of *0* to yield consistently positive signs for *ν* estimates. BAGEA was run for 300 iterations in each analysis.

## 5.6 Author Contributions

D.L. implemented the software and performed experiments. D.L, R.B, K.H., G.B. V.H.S. wrote the manuscript.

## 5.8 Acknowledgements

Special thanks to Prof. Zoltan Kutalik for helpful discussions.

## 5.9 Download Links

- UCSC: http://hgdownload.cse.ucsc.edu/goldenpath/hg19/database/
- EXPECTO: https://github.com/FunctionLab/ExPecto/
- 1KG: ftp://ftp-trace.ncbi.nih.gov/1000genomes/ftp/release/20130502/
- GTEX: https://gtexportal.org/home/datasets
- GEUVADIS: ftp://ftp.ebi.ac.uk/pub/databases/microarray/data/experiment/GEUV/E-GEUV-3/analysis_results/
- BAGEA: https://github.com/dlampart/bagea

