## Supplementary Information for "A Framework for Integrating Directed and Undirected Annotations to Build Explanatory Models of cis-eQTL Data"

### 1 Supplementary Information

#### 1.1 Outline of Variational Message Passing

To fit the *BAGEA* model, we use variational message passing[1]. We will very briefly outline the algorithm but encourage the reader to consult the literature for a fuller understanding. The model prescribes a distribution $P[\boldsymbol{H}, \boldsymbol{V}]$ over the hidden nodes $\boldsymbol{H}$ and visible nodes $\boldsymbol{V}$. The goal is to be able to marginalize over this model such that we can compute, say, $E[\boldsymbol{H_i}|\boldsymbol{V}]$ for a particular node $\boldsymbol{H_i}$ of interest. Variational message passing approximates $P[\boldsymbol{H}|\boldsymbol{V}]$ with a more tractable distribution $Q(\boldsymbol{H})$ over the hidden nodes with the property that each hidden node in the network is independent of all others (i.e. $Q(\boldsymbol{H}) = \prod_i Q_{H_i}(\boldsymbol{H_i})$). This distribution is chosen such that it minimizes the Kullback-Leibler (KL) divergence to $P[\boldsymbol{H}|\boldsymbol{V}]$. Variational message passing is designed to iteratively update $Q$ to minimize the KL divergence (convergence to the optimum is not guaranteed however, as the procedure can get trapped in local optima). It can be regarded as coordinate descent, where the distribution $Q_{H_i}$ gets updated separately while keeping the other distributions $Q_{H_j}$ $j \neq i$ constant. It can be shown that the optimal distribution $Q^*_{H_i}$ in terms of the KL divergence has the following property: $\log(Q^*_{H_i}(\boldsymbol{H_i})) =< \log(P[\boldsymbol{V}, \boldsymbol{H}]) >_{\sim Q_{H_i}(\boldsymbol{H_i})}$, where $< . >_{\sim Q_{H_i}(\boldsymbol{H_i})}$ refers to the expectation w.r.t to all $Q_{H_j}(\boldsymbol{H_j})$ with $i \neq j$.

##### 1.1.1 Conjugate-Exponential Models

To be able to run variational message passing, the model has to be conjugate-exponential. To satisfy the exponentiality condition, all random variable nodes, conditional on their parent nodes, have to be members of the exponential family and should be writable in the following form:

$$\log(P[\boldsymbol{x}]) = \boldsymbol{\phi_x}^T \boldsymbol{u_x}(\boldsymbol{x}) + g(\boldsymbol{\phi_x}) + f(\boldsymbol{x}), \quad (1)$$

where $\boldsymbol{u_x}(\boldsymbol{x})$ is the sufficient statistic vector and $\boldsymbol{\phi_x}$ is the natural parameter vector. To make the conditional character explicit, we can write

$$\log(P[\boldsymbol{x}|\boldsymbol{pa_x}]) = \boldsymbol{\phi_x}(\boldsymbol{pa_x})^T \boldsymbol{u_x}(\boldsymbol{x}) + g(\boldsymbol{\phi_x}(\boldsymbol{pa_x})) + f(\boldsymbol{x}), \tag{2}$$

where $\boldsymbol{pa_x}$ denotes all parents of $\boldsymbol{x}$. To satisfy the conjugacy condtion, the conditional log probability of $\boldsymbol{x}$ needs to follow the same functional form as the log probability of its parent $\boldsymbol{y}$:

$$\log(P[\boldsymbol{x}|\boldsymbol{y}, \boldsymbol{cp_y(x)}]) = \boldsymbol{\phi_{xy}(x, cp_y(x))}^T \boldsymbol{u_y}(\boldsymbol{y}) + \lambda(\boldsymbol{x}, \boldsymbol{cp_y(x)}), \tag{3}$$

where $\boldsymbol{cp_y(x)}$ denotes all other parents of $\boldsymbol{x}$ except $\boldsymbol{y}$, i.e. the coparents of $\boldsymbol{y}$. Put another way, one needs to be able to express the dependence of $\log(P[\boldsymbol{x}|\boldsymbol{y}, \boldsymbol{cp_y(x)}])$ on $\boldsymbol{y}$ as a linear function of the sufficient statistic vector $\boldsymbol{u_y}(\boldsymbol{y})$. Analogously, this condition has to hold for all other parents of $\boldsymbol{x}$ too. As a consequence of this, we can write the entire distribution $\log(P[\boldsymbol{V}, \boldsymbol{H}])$ as depending on $\boldsymbol{y}$ via its sufficient statistic vector $\boldsymbol{u_y}(\boldsymbol{y})$ as follows:

$$\log(P[\boldsymbol{V}, \boldsymbol{H}]) = \Big(\boldsymbol{\phi_y(pa_y)} + \sum_{x_i \in \boldsymbol{ch_y}} \boldsymbol{\phi_{x_iy}(x_i, cp_y(x_i))}\Big)\boldsymbol{u_y}(\boldsymbol{y}) + f(\boldsymbol{y}) + \boldsymbol{g}, \tag{4}$$

where $\boldsymbol{g}$ is independent of $\boldsymbol{y}$, $\boldsymbol{ch_y}$ denotes the set of children of $\boldsymbol{y}$ and $\boldsymbol{cp_y(x_i))}$ denotes the set of parents of $\boldsymbol{x_i}$ except $\boldsymbol{y}$. As we can see, if we fix all variables except $\boldsymbol{y}$, the resulting distribution is in the same exponential family as $\boldsymbol{x}$ conditional on its parents, i.e. $\boldsymbol{x}|\boldsymbol{pa_x}$. Moreover, if we take expectations w.r.t. to $< . >_{\sim Q_y(y)}$, we see that the resulting distribution still is in the same exponential family. To put it another way: for any $\boldsymbol{H_i}$, $Q_{H_i}(\boldsymbol{H_i})$ has the same functional form as $P[\boldsymbol{H_i}|\boldsymbol{pa_{H_i}}]$, where $\boldsymbol{pa_{H_i}}$ is the parents of $\boldsymbol{H_i}$. For instance, if $P[\boldsymbol{H_i}|\boldsymbol{pa_{H_i}}]$ has a normal density, then so does $Q_i(\boldsymbol{H_i})$.

To update $Q_{\boldsymbol{y}}(\boldsymbol{y})$, we need to be able to calculate $< \boldsymbol{\phi_y}(\boldsymbol{pa_y}) >_{\sim Q_{\boldsymbol{y}}}$ and $\boldsymbol{\phi_{x_iy}}(\boldsymbol{x_i}, \boldsymbol{cp_y(x_i)})_{\sim Q_{\boldsymbol{y}}}$ for all children $\boldsymbol{x_i}$ of $\boldsymbol{y}$. This will allow us to calculate the natural parameter vector for the updated $Q_{\boldsymbol{y}}$. Note that, because of conjugacy, and all $Q_{H_i}$ being independent from each other, $< \boldsymbol{\phi_{xy}(x, cp_y)} >$ can be expressed as a function of the expectation of the sufficient statistic vector $E_{Q_{\boldsymbol{x}}}[\boldsymbol{u_x}(x)]$ as well as the expectation of the sufficient statistic vector of all coparents of $\boldsymbol{y}$.

##### 1.1.2 Form of Messages

The message passing algorithm works by sending messages to neighboring nodes and local updates at the nodes once the messages are gathered. Messages sent from child $\boldsymbol{x}$ to parent $\boldsymbol{y}$ has the form

$$< \boldsymbol{\phi_{xy}(x, cp_y(x))} >, \tag{5}$$

where $< \cdot >$ denotes the expectation with respect $Q(\boldsymbol{H})$ (in practice we do not know $Q(\boldsymbol{H})$ as this is the quantity we are trying to find. We plug in our current estimate instead). Again, because of conjugacy and all $Q_{H_i}$ being independent from each other, $< \boldsymbol{\phi_{xy}(x, cp_y)} >$ can be expressed as a function of the expectation of the sufficient statistic vector $E[\boldsymbol{u_x}]$ as well as the expectation of the sufficient statistic vector of all coparents of $\boldsymbol{y}$. Messages passed from a parent $\boldsymbol{z}$ of $\boldsymbol{y}$ to $\boldsymbol{y}$ is just the expectation of the sufficient statistic vector $E[\boldsymbol{u_z}]$ itself. These messages are then used to compute

$$< \boldsymbol{\phi_y(pa_y)} > \tag{6}$$

Once the messages are received, we want to update $Q_{\boldsymbol{y}}(\boldsymbol{y})$ itself. As we noted before, $Q_{\boldsymbol{y}}(\boldsymbol{y})$ has the same functional form as $P[\boldsymbol{y}|\boldsymbol{pa_y}]$, i.e. it is in the same exponential family. We therefore only need to find its natural parameter vector $\boldsymbol{\phi_y^*}$. This can be computed from the sent messages:

$$\boldsymbol{\phi_y^*} =< \boldsymbol{\phi_y(pa_z)} > + \sum_{x_i \in \boldsymbol{ch_y}} < \boldsymbol{\phi_{x_i y}(x_i, cp_y(x_i))} >, \tag{7}$$

where $\boldsymbol{ch_y}$ denotes the set of children of $\boldsymbol{y}$ and $\boldsymbol{cp_y(x_i)})$ denotes the set of parents of $\boldsymbol{x_i}$ except $\boldsymbol{y}$. Once we know $\boldsymbol{\phi_y^*}$, we have implicitly determined an updated distribution $Q_{\boldsymbol{y}}(\boldsymbol{y})$ and can find the coresponding expectations of its sufficient statistic vector $E[\boldsymbol{u_y}]$. This serves again as the message to all children of $\boldsymbol{y}$ and is used in the computation of messages to its parents.

#### 1.2 Implementing the *BAGEA* Model

##### 1.2.1 Log-Probability Densities

In the following, we list all log probability densities of the *BAGEA* model described in the methods section:

$$\log(P[\boldsymbol{y}_j|\lambda_j,\boldsymbol{b}_j]) = -\frac{n}{2}\log(2\pi)+\frac{n}{2}\log(\lambda_j)-\frac{\lambda_j}{2}(\boldsymbol{y}_j^T\boldsymbol{y}_j-2\boldsymbol{y}_j^T\boldsymbol{X}_j\boldsymbol{b}_j+\boldsymbol{b}_j^T\boldsymbol{X}_j^T\boldsymbol{X}_j\boldsymbol{b}_j),$$

$$\log(P[\lambda_j|\lambda_1,\lambda_2]) = \lambda_1\log(\lambda_2)-\log(\Gamma(\lambda_1))+(\lambda_1-1)\log(\lambda_j)-\lambda_2\lambda_j,$$

$$\log(P[\lambda_2|\rho_1,\rho_2]) = \rho_1\log(\rho_2)-\log(\Gamma(\rho_1))+(\rho_1-1)\log(\lambda_2)-\rho_2\lambda_2,$$

$$\log(P[\alpha_{ij}|\gamma_1,\gamma_{ij},\kappa_j]) = \gamma_1\Big(\log(\kappa_j)+\log(\gamma_{ij})\Big)-\log(\Gamma(\gamma_1))+(\gamma_1-1)\log(\alpha_{ij})-\kappa_j\gamma_{ij}\alpha_{ij},$$

i.e.:

$$\log(P[\alpha_{ij}|a_1,..,a_t]) = \gamma_1\Big(\log(\kappa_j)+\sum_{k:\boldsymbol{C}^{\boldsymbol{j}}_{ik}=1}\log(a_k)\Big)-\log(\Gamma(\gamma_1))+(\gamma_1-1)\log(\alpha_{ij})-\alpha_{ij}\kappa_j\prod_{k:\boldsymbol{C}^{\boldsymbol{j}}_{ik}=1}a_k,$$

$$\log(P[a_k|\phi_1,\phi_2]) = \phi_1\log(\phi_2)-\log(\Gamma(\phi_1))+(\phi_1-1)\log(a_k)-\phi_2 a_k,$$

$$\log(P[\kappa_j|\tau_1,\tau_2]) = \tau_1\log(\tau_2)-\log(\Gamma(\tau_1))+(\tau_1-1)\log(\kappa_j)-\tau_2\kappa_j,$$

$$\log(P[\tau_2|\xi_1,\xi_2]) = \xi_1\log(\xi_2)-\log(\Gamma(\xi_1))+(\xi_1-1)\log(\tau_2)-\xi_2\tau_2,$$

$$\begin{aligned}\log(P[b_{ij}|\boldsymbol{\nu},\boldsymbol{\omega},\alpha_{ij}]) = &-\frac{1}{2}\log(2\pi)+\frac{1}{2}\log(\alpha_{ij})-\frac{\alpha_{ij}}{2}(b_{ij}^2+((\boldsymbol{v}_{\boldsymbol{i}}^{\boldsymbol{j}})^T\boldsymbol{\omega})^2((\boldsymbol{f}_{\boldsymbol{i}}^{\boldsymbol{j}})^T\boldsymbol{\nu})^2\\ &-2b_{ij}((\boldsymbol{v}_{\boldsymbol{i}}^{\boldsymbol{j}})^T\boldsymbol{\omega})((\boldsymbol{f}_{\boldsymbol{i}}^{\boldsymbol{j}})^T\boldsymbol{\nu})),\end{aligned}$$

where $\boldsymbol{v}_{\boldsymbol{i}}^{\boldsymbol{j}}$ and $\boldsymbol{f}_{\boldsymbol{i}}^{\boldsymbol{j}}$ are the $i$th row vector of $\boldsymbol{V}^{\boldsymbol{j}}$ and $\boldsymbol{F}^{\boldsymbol{j}}$ respectively. Further,

$$\log(P[\boldsymbol{\nu}]) = -\frac{q}{2}\log(2\pi)+\frac{1}{2}\sum_{i=1}^{q}\log(p_i)-\sum_{i=1}^{q}\frac{p_i}{2}(\nu_i^2+c_i^2-2c_i\nu_i),$$

$$\log(P[\omega_i|\boldsymbol{v}^{\boldsymbol{1}},..,\boldsymbol{v}^{\boldsymbol{q}}]) = \frac{-1}{2}\log(2\pi)+\frac{1}{2}(\sum_{j=1}^{q}\log(v^j_{\boldsymbol{d}_i^j}))-\prod_{j=1}^{q}v^j_{\boldsymbol{d}_i^j}\frac{\omega_i^2}{2},$$

$$\log(P[\upsilon_k^j|\chi_{1j},\chi_{2j}]) = \chi_{1j}\log(\chi_{2j}) - \log(\Gamma(\chi_{1j})) + (\chi_{1j}-1)\log(\upsilon_k^j) - \chi_{2j}\upsilon_k^j,$$

$$\log(P[\chi_{2j}|\zeta_1,\zeta_2]) = \zeta_1\log(\zeta_2) - \log(\Gamma(\zeta_1)) + (\zeta_1-1)\log(\chi_{2j}) - \zeta_2\chi_{2j}.$$

##### 1.2.2 Expectation of Natural Parameter Vector $\phi$: w.r.t $Q$

$$< \phi_{b_{ij}} >= \Big(E_u[\boldsymbol{u}_{\alpha_{ij}}(2)](\boldsymbol{f_i^j})^T E_u[\boldsymbol{u_\nu}(1)](\boldsymbol{v_i^j})^T E_u[\boldsymbol{u_\omega}(1)], -\frac{E_u[\boldsymbol{u}_{\alpha_{ij}}(2)]}{2}\Big),$$

$$< \phi_{\boldsymbol{\omega}} >= (\boldsymbol{0}^T, vec(-\frac{1}{2}diag(< \boldsymbol{\delta} >))^T)^T,$$

$$< \delta_i >= \prod_{j=1}^{q} E[\boldsymbol{u}_{\upsilon^j_{\boldsymbol{d}_i^j}}(2)],$$

$$< \boldsymbol{\phi}_{\upsilon_k^j} >= ((\chi_{1j}-1), -E_u[\boldsymbol{u}_{\chi_{2j}}(2)]),$$

$$< \boldsymbol{\phi}_{\chi_{2j}} >= ((\zeta_1-1), -\zeta_2)^T,$$

$$< \phi_{\boldsymbol{\nu}} >= \Big((\boldsymbol{c}\cdot\boldsymbol{p})^T, -\frac{1}{2}diag(\boldsymbol{p})\Big),$$

$$< \boldsymbol{\phi}_{\alpha_{ij}} >= (\gamma_1 - 1, -E[\boldsymbol{u}_{\kappa_j}(2)] \prod_{k:\boldsymbol{C}_{ik}^{\boldsymbol{j}}=1} E_u[\boldsymbol{u}_{a_k}(2)]),$$

$$< \boldsymbol{\phi}_{a_k} >= ((\phi_1-1), -\phi_2),$$

$$< \boldsymbol{\phi}_{\lambda_j} >= ((\lambda_1-1), -E_u[\boldsymbol{u}_{\lambda_2}(2)]),$$

$$< \boldsymbol{\phi}_{\lambda_2} >= ((\rho_1-1), -\rho_2),$$

$$< \boldsymbol{\phi}_{\kappa_j} >= ((\tau_1-1), -E_u[\boldsymbol{u}_{\tau_2}(2)]),$$

$$< \boldsymbol{\phi}_{\tau_2} >= ((\xi_1-1), -\xi_2).$$

##### 1.2.3 Expectation of Sufficient Statistic Vector $E[\boldsymbol{u_x}]$ as a Function of $\phi^*_{\boldsymbol{x}}$ (Messages to Children)

$$E_u[\boldsymbol{u}_{a_k}] = \Big(\psi(\phi^*_{a_k}(1)+1) - \log(-\phi^*_{a_k}(2)), \frac{-\phi^*_{a_k}(1)-1}{\phi^*_{a_k}(2)}\Big),$$

$$E_u[\boldsymbol{u}_{\alpha_{ij}}] = \Big(\psi(\phi^*_{\alpha_{ij}}(1)+1) - \log(-\phi^*_{\alpha_{ij}}(2)), \frac{-\phi^*_{\alpha_{ij}}(1)-1}{\phi^*_{\alpha_{ij}}(2)}\Big),$$

$$E[\boldsymbol{u}_{\lambda_j}] = \Big(\psi(\phi^*_{\lambda_j}(1)+1) - \log(-\phi^*_{\lambda_j}(2)), \frac{-\phi^*_{\lambda_j}(1)-1}{\phi^*_{\lambda_j}(2)}\Big),$$

$$E[\boldsymbol{u}_{\lambda_2}] = \Big(\psi(\phi^*_{\lambda_2}(1)+1) - \log(-\phi^*_{\lambda_2}(2)), \frac{-\phi^*_{\lambda_2}(1)-1}{\phi^*_{\lambda_2}(2)}\Big),$$

$$E[\boldsymbol{u}_{\kappa_j}] = \Big(\psi(\phi^*_{\kappa_j}(1)+1) - \log(-\phi^*_{\kappa_j}(2)), \frac{-\phi^*_{\kappa_j}(1)-1}{\phi^*_{\kappa_j}(2)}\Big),$$

$$E[\boldsymbol{u}_{\tau_2}] = \Big(\psi(\phi^*_{\tau_2}(1)+1) - \log(-\phi^*_{\tau_2}(2)), \frac{-\phi^*_{\tau_2}(1)-1}{\phi^*_{\tau_2}(2)}\Big),$$

$$E[\boldsymbol{u}_{\boldsymbol{b}_j}] = (E[\boldsymbol{u}_{\boldsymbol{b}_j}(1)], E[\boldsymbol{u}_{\boldsymbol{b}_j}(2)]),$$

$$E[\boldsymbol{u}_{\boldsymbol{b}_j}] = (-\boldsymbol{\phi}^*_{\boldsymbol{b_j}}(2)^{-1}\boldsymbol{\phi}^*_{\boldsymbol{b_j}}(1)/2, vec(-\boldsymbol{\phi}^*_{\boldsymbol{b_j}}(2)^{-1}/2 + E[\boldsymbol{u}_{\boldsymbol{b_j}}(1)]E[\boldsymbol{u}_{\boldsymbol{b_j}}(1)]^T)),$$

$$E[\boldsymbol{u}_{\omega}] = (-\boldsymbol{\phi}^*_{\boldsymbol{\omega}}(2)^{-1}\boldsymbol{\phi}^*_{\boldsymbol{\omega}}(1)/2, vec(-\boldsymbol{\phi}^*_{\boldsymbol{\omega}}(2)^{-1}/2 + E[\boldsymbol{u}_{\boldsymbol{\omega}}(1)]E[\boldsymbol{u}_{\boldsymbol{\omega}}(1)]^T)),$$

$$E_u[\boldsymbol{u}_{\chi_{2j}}] = \Big(\psi(\phi^*_{\chi_{2j}}(1)+1) - \log(-\phi^*_{\chi_{2j}}(2)), \frac{-\phi^*_{\chi_{2j}}(1)-1}{\phi^*_{\chi_{2j}}(2)}\Big),$$

$$E_u[\boldsymbol{u}_{\upsilon_k^j}] = \Big(\psi(\phi^*_{\upsilon_k^j}(1)+1) - \log(-\phi^*_{\upsilon_k^j}(2)), \frac{-\phi^*_{\upsilon_k^j}(1)-1}{\phi^*_{\upsilon_k^j}(2)}\Big),$$

$$E[\boldsymbol{u}_{\nu}] = (-\boldsymbol{\phi}^*_{\boldsymbol{\nu}}(2)^{-1}\boldsymbol{\phi}^*_{\boldsymbol{\nu}}(1)/2, vec(-\boldsymbol{\phi}^*_{\boldsymbol{\nu}}(2)^{-1}/2 + E[\boldsymbol{u}_{\boldsymbol{\nu}}(1)]E[\boldsymbol{u}_{\boldsymbol{\nu}}(1)]^T)).$$

##### 1.2.4 Messages from Children to Parents

$$<\boldsymbol{\phi}_{\alpha_{ij}a'_k}> = \Big(\gamma_1 \mathbf{1}_{\{k' \in C_{ij}\}}, -E[\boldsymbol{u}_{\kappa_j}(2)]\Big(\prod_{k \neq k' \in C_{ij}} E_u[\boldsymbol{u}_{a_k}(2)]\Big)E_u[\boldsymbol{u}_{\alpha_{ij}}(2)]\Big),$$

$$<\boldsymbol{\phi}_{\alpha_{ij}\kappa_j}> = \Big(\gamma_1, -(\prod_{k \in C_{ij}} E_u[\boldsymbol{u}_{a_k}(2)])E_u[\boldsymbol{u}_{\alpha_{ij}}(2)]\Big),$$

$$<\boldsymbol{\phi}_{\boldsymbol{y}_j\lambda_j}> \;=\; (\frac{n}{2}, \frac{-1}{2}(\boldsymbol{y}_j^T\boldsymbol{y}_j - 2\boldsymbol{y}_j^T\boldsymbol{X}_jE_u[\boldsymbol{u}_{\boldsymbol{b_j}}(1)] + Tr(\boldsymbol{X}_j^T\boldsymbol{X}_jE_u[\boldsymbol{u}_{\boldsymbol{b_j}}(2)])))^T,$$

$$<\boldsymbol{\phi}_{\boldsymbol{y}_j\boldsymbol{b}_j}> \;=\; ((E[\boldsymbol{u}_{\lambda_j}]\boldsymbol{y}_j^T\boldsymbol{X}_j)^T, -(E[\boldsymbol{u}_{\lambda_j}]vec(\boldsymbol{X}_j^T\boldsymbol{X}_j)/2)^T)^T,$$

$$< \boldsymbol{\phi}_{\lambda_j \lambda_2} > \; = \; (\lambda_1, -E_u[\boldsymbol{u}_{\lambda_j}(2)]),$$

$$< \boldsymbol{\phi}_{\kappa_j \tau_2} > \; = \; (\tau_1, -E_u[\boldsymbol{u}_{\kappa_j}(2)]),$$

$$< \phi_{b_{ij}\boldsymbol{\alpha_{ij}}} >= \Big(\frac{1}{2}, -\frac{1}{2}(E_u[\boldsymbol{u}_{\boldsymbol{b_j}}(2)]_{ii} + (\boldsymbol{f_{ij}}^T E_u[\boldsymbol{u}_\nu(2)]\boldsymbol{f_{ij}}\boldsymbol{v_{ij}}^T E_u[\boldsymbol{u}_\omega(2)]\boldsymbol{v_{ij}}, - 2E_u[\boldsymbol{u}_{\boldsymbol{b_j}}(1)]_i \boldsymbol{v_{ij}}^T E_u[\boldsymbol{u}_\omega(1)]\boldsymbol{f_{ij}}^T E_u[\boldsymbol{u}_\nu(1)]))\Big),$$

$$< \phi_{b_{ij}\boldsymbol{\omega}} >= \Big(E_u[\boldsymbol{u}_{\alpha_{ij}}(2)]E_u[\boldsymbol{u}_{\boldsymbol{b_j}}(1)]_i \boldsymbol{f_{ij}}^T E_u[\boldsymbol{u}_\nu(1)]\boldsymbol{v_{ij}}^T, (-\frac{E_u[\boldsymbol{u}_{\alpha_{ij}}(2)]\boldsymbol{f_{ij}}^T E_u[\boldsymbol{u}_\nu(2)]\boldsymbol{f_{ij}}}{2} vec(\boldsymbol{v_{ij}}\boldsymbol{v_{ij}}^T))^T\Big),$$

$$< \phi_{b_{ij}\boldsymbol{\nu}} >= \Big(E_u[\boldsymbol{u}_{\alpha_{ij}}(2)]E_u[\boldsymbol{u}_{\boldsymbol{b_j}}(1)]_i \boldsymbol{v_{ij}}^T E_u[\boldsymbol{u}_\omega(1)]\boldsymbol{f_{ij}}^T, (-\frac{E_u[\boldsymbol{u}_{\alpha_{ij}}(2)]\boldsymbol{v_{ij}}^T E_u[\boldsymbol{u}_\omega(2)]\boldsymbol{v_{ij}}}{2} vec(\boldsymbol{f_{ij}}\boldsymbol{f_{ij}}^T))^T\Big),$$

$$< \boldsymbol{\phi}_{\omega_i \upsilon_k^j} >= \mathbf{1}_{\{k=\boldsymbol{d}_i^j\}}\Big(\frac{1}{2}, -\Big(\prod_{j' \neq j} E[\boldsymbol{u}_{\upsilon^{j'}_{\boldsymbol{d}_i^{j'}}}(2)]\Big)\frac{E[\boldsymbol{u}_\omega(2)]_{ii}}{2}\Big),$$

$$< \boldsymbol{\phi}_{\upsilon_k^j \chi_{2j}} > \; = \; (\chi_{1j}, -E_u[\upsilon_k^j(2)]).$$

##### 1.2.5 Approximating -$\phi_{\boldsymbol{b}}^*(2)$ for Fast Inversion

One time critical step when performing the updates is the inversion of $\boldsymbol{\phi}_{\boldsymbol{b}}^*(2)$ which has to be done for each round and for each gene.

$$\begin{aligned} -\boldsymbol{\phi}_{\boldsymbol{b}}^*(2) &= E[\boldsymbol{u}_\lambda(2)](\boldsymbol{X}^T\boldsymbol{X})/2) + \frac{1}{2}diag(E[\boldsymbol{u}_{\alpha_i}(2)]), \\ &= c\boldsymbol{X}^T\boldsymbol{X} + \boldsymbol{D}_{\boldsymbol{\alpha}}, \end{aligned}$$

where $c$ is a scalar and $\boldsymbol{D}_{\boldsymbol{\alpha}}$ is a diagonal matrix (gene-wise indices were dropped). Because of LD and data on fewer samples than SNPs in the gene region, $\boldsymbol{X}^T\boldsymbol{X}$ is degenerate or close to degeneracy and we can approximate it with a low rank matrix. Set $\boldsymbol{X} = \boldsymbol{U}\boldsymbol{D}\boldsymbol{V}^T$ as the singular value decomposition. Then, set $\boldsymbol{U_t}$ to consist of the first $t$ columns and $\boldsymbol{D_t}$ to consist of the upper left $t \times t$ submatrix of $\boldsymbol{D}$. $t$ is set such that the sum of squares of $\boldsymbol{D_t}$ is close to the sum of squares of $\boldsymbol{D}$ (say 99%). Set $\boldsymbol{A_t} = \sqrt{c}\boldsymbol{U_t D_t}$ so that

$$c\boldsymbol{X^T X} + \boldsymbol{D_\alpha} \quad \approx \quad \boldsymbol{A}_t \boldsymbol{A}_t^T + \boldsymbol{D_\alpha}.$$

Using the Woodbury matrix identity, we have

$$(\boldsymbol{A}_t \boldsymbol{A}_t^T + \boldsymbol{D_\alpha})^{-1} \quad = \quad \boldsymbol{D_\alpha}^{-1} - \boldsymbol{D_\alpha}^{-1} \boldsymbol{A}_t (\boldsymbol{I}_t + \boldsymbol{A}_t^T \boldsymbol{D_\alpha}^{-1} \boldsymbol{A}_t)^{-1} \boldsymbol{A}_t^T \boldsymbol{D_\alpha}^{-1}.$$

Since $\boldsymbol{D_\alpha}$ is diagonal it is easy to invert and $(\boldsymbol{I_t} + \boldsymbol{A}_t^T \boldsymbol{D_\alpha}^{-1} \boldsymbol{A}_t)$ is of dimension $t$ and can be inverted much faster than the full matrix. This approach builds on the fact that the matrix to invert is well conditioned if $\boldsymbol{D_\alpha}$ is large enough (relative to the diagonal of $c\boldsymbol{X}^T\boldsymbol{X}$), i.e.: if the regression is regularized sufficiently via the priors on $\boldsymbol{b}$ and $\lambda$. We can also add the discarded variance onto the diagonal. Set $t^c$ as the indices larger than $t$ and $\boldsymbol{A_{t^c}} = \sqrt{c}\boldsymbol{U_{t^c} D_{t^c}}$. Then $w = \sum_{ij}([\boldsymbol{A_{t^c}}]_{ij})^2$ is the the total variance discarded. We set $\boldsymbol{D'_\alpha} = \boldsymbol{D_\alpha} + \boldsymbol{I}_n \frac{w}{n}$, where $n$ is the dimension of $\boldsymbol{D_\alpha}$

$$(c\boldsymbol{X}^T\boldsymbol{X} + \boldsymbol{D_\alpha})^{-1} \quad \approx \quad \boldsymbol{D'_\alpha}^{-1} - \boldsymbol{D'_\alpha}^{-1} \boldsymbol{A}_t (\boldsymbol{I}_t + \boldsymbol{A}_t^T \boldsymbol{D'_\alpha}^{-1} \boldsymbol{A}_t)^{-1} \boldsymbol{A}_t^T \boldsymbol{D'^{-1}_\alpha}.$$

Additionally to the inversion, we would like to speed up the determinant calculation step used to calculate the lower bound. We can make use of the matrix determinant lemma i.e.

$$\det(\boldsymbol{D'}_\alpha + \boldsymbol{A}_t \boldsymbol{A}_t^T) = \det(\boldsymbol{I_t} + \boldsymbol{A}_t^T \boldsymbol{D'^{-1}_\alpha} \boldsymbol{A}_t) \det(\boldsymbol{D'_\alpha}).$$

##### 1.2.6 Working with Summary Statistics.

As we mentioned above, we can approximate $\boldsymbol{X}^T\boldsymbol{X}$. We can also estimate it externally from 1000 Genomes, with $\boldsymbol{\Sigma}$ say, and approximate $\boldsymbol{X^T y}$ with summary statistics $\sqrt{n}\boldsymbol{z}$. However, this can lead to convergence problems as the modeling assumptions do not hold anymore. For limited $n$ and arbitrary but fixed $\boldsymbol{X}^T\boldsymbol{X}$ and $\boldsymbol{X^T y}$, there is not always an adequate $\boldsymbol{X}$ and $\boldsymbol{y}$ fullfilling the equations. We therefore reformulate the model in terms of summary statistics such that for each SNP-gene pair the statistics $\boldsymbol{z}_j$ are observed: Since we have

$$\frac{1}{\sqrt{n}}\boldsymbol{X}_j^T\boldsymbol{y}_j = \frac{1}{\sqrt{n}}\boldsymbol{X}_j^T\boldsymbol{X}_j\boldsymbol{b}_j + \frac{1}{\sqrt{n}}\boldsymbol{X}_j^T\boldsymbol{\epsilon}_j.$$

We can naturally model the vector of summary statistics as

$$\boldsymbol{z}_j = \boldsymbol{\Sigma}_j\sqrt{n}\boldsymbol{b}_j + \boldsymbol{\epsilon}'_j,$$

where $\boldsymbol{\epsilon}'_j \sim N_m(\boldsymbol{0}, \lambda_j^{-1}\boldsymbol{\Sigma}_j)$.

We then have

$$\log(P[\boldsymbol{z}_j|\boldsymbol{b}_j]) = -\frac{m_j}{2}\log(2\pi) - \frac{1}{2}\log(\det(\frac{1}{\lambda_j}\boldsymbol{\Sigma}_j)) - \frac{\lambda_j}{2}(\boldsymbol{z}_j^T\boldsymbol{\Sigma}_j^{-1}\boldsymbol{z}_j - 2\sqrt{n}\boldsymbol{z}_j^T\boldsymbol{b}_j + n\boldsymbol{b}'^T_j\boldsymbol{\Sigma}_j\boldsymbol{b}_j).$$

If $\boldsymbol{\Sigma}_j$ is not full rank we could use the pseudo-inverse. Note however, that in practice, regularization is needed as otherwise $E[\boldsymbol{z}]$ will most likely not lie in the vector space spanned by the eigenvectors of $\boldsymbol{\Sigma}_j$.If $\boldsymbol{\Sigma_j}$ has rank $n$, we have,

$$\boldsymbol{\phi}_{\boldsymbol{z}_j\boldsymbol{b}_j} \quad = \quad ((\sqrt{n}\lambda_j\boldsymbol{z}_j), -(n\lambda_j vec(\boldsymbol{\Sigma}_j)/2)^T)^T.$$

Next, we have

$$\boldsymbol{\phi}_{\boldsymbol{z}_j\lambda_j} \quad = \quad (\frac{m_j}{2}), -\frac{1}{2}(\boldsymbol{z}_j^T\boldsymbol{\Sigma}_j^{-1}\boldsymbol{z}_j - 2\sqrt{n}\boldsymbol{z}_j^T\boldsymbol{b}_j + n\boldsymbol{b}_j{}^T\boldsymbol{\Sigma}_j\boldsymbol{b}_j).$$

Importantly, if $\boldsymbol{\Sigma}_j$ is full rank, the right hand term is now guaranteed to be non-positive which could not be guaranteed before the reformulation.

##### 1.2.7 Computing the Lower Bound of the Data Log-Evidence

Variational message passing can additionally provide a lower bound $L(Q)$ on the data log-evidence $\log(P[V])$, where the gap is exactly the KL divergence. To compute the $L(Q)$ the algorithm needs to additionally compute $< g(\boldsymbol{\phi}_{\boldsymbol{x}}(\boldsymbol{pa}_{\boldsymbol{x}})) >$, $g(\boldsymbol{\phi^*}_{\boldsymbol{x}})$ and $\log(P[\boldsymbol{V}|\boldsymbol{H}])$. For the *BAGEA* model, the required quantities of the form $< g(\boldsymbol{\phi_x}(\boldsymbol{pa_x})) >$ are:

$$< g_{b_{ij}}(pa) >= \frac{1}{2}E[\boldsymbol{u}_{\alpha_{ij}}(1)] - \frac{E[\boldsymbol{u}_{\alpha_{ij}}(2)]}{2}Tr(\boldsymbol{v_{ij}}\boldsymbol{v_{ij}}^T E[\boldsymbol{u_\omega}(2)])Tr(\boldsymbol{f_{ij}}\boldsymbol{f_{ij}}^T E[\boldsymbol{u_\nu}(2)]),$$

$$< g_{\lambda_j}(pa) >= \lambda_1 E[\boldsymbol{u}_{\lambda_2}(1)] - \log(\Gamma(\lambda_1)),$$

$$< g_{\lambda_2}(pa) >= \rho_1 \log(\rho_2) - \log(\Gamma(\rho_1)),$$

$$< g_{\alpha_i}(pa) >= \gamma_1 \Big(E[\boldsymbol{u}_{\kappa_j}(1)] + \sum_{k:\boldsymbol{C}^{\boldsymbol{j}}_{ik}=1} E[\boldsymbol{u}_{a_k}(1)]\Big) - \log(\Gamma(\gamma_1)),$$

$$< g_{\kappa_j}(pa) >= \tau_1 E[\boldsymbol{u}_{\tau_2}(1)] - \log(\Gamma(\tau_1)),$$

$$< g_{\tau_2}(pa) >= \xi_1 \log(\xi_2) - \log(\Gamma(\xi_1)),$$

$$< g_{a_k}(pa) >= \phi_1 \log(\phi_2) - \log(\Gamma(\phi_1)),$$

$$< g_{\omega_i}(pa) >= \frac{-1}{2}\log(2\pi) + \frac{1}{2}(\sum_{j=1}^{q} E[\boldsymbol{u}_{\upsilon^j_{\boldsymbol{d}^j_i}}(1)]),$$

$$< g_{\upsilon^j_k}(pa) >= \chi_{1j} E_u[\boldsymbol{u}_{\chi_{2j}}(1)] - \log(\Gamma(\chi_{1j})),$$

$$< g_{\boldsymbol{\nu}}(pa) >= \frac{-q}{2}\log(2\pi) + \frac{1}{2}\sum_{i=1}^{q}\log(p_i) - \sum_{i=1}^{q}\frac{p_i c_i^2}{2},$$

$$< g_{\chi_{wj}}(pa) >= \zeta_1 \log(\zeta_2) - \log(\Gamma(\zeta_1)).$$

the required quantities of the form $g(\boldsymbol{\phi^*_x})$ are:

$$g(\boldsymbol{\phi}^*_{\boldsymbol{b}_j}) = -\frac{m_j}{2}\log(2\pi) + \frac{1}{2}\log(\det(-2(\boldsymbol{\phi}^*_{\boldsymbol{b}_j}(2)))) + \frac{1}{4}(\boldsymbol{\phi}^*_{\boldsymbol{b}_j}(1))(\boldsymbol{\phi}^*_{\boldsymbol{b}_j}(2))^{-1}(\boldsymbol{\phi}^*_{\boldsymbol{b}_j}(1))^T,$$

$$g(\boldsymbol{\phi}^*_{\boldsymbol{\omega}}) = -\frac{s}{2}\log(2\pi) + \frac{1}{2}\log(\det(-2(\boldsymbol{\phi}^*_{\boldsymbol{\omega}}(2)))) + \frac{1}{4}(\boldsymbol{\phi}^*_{\boldsymbol{\omega}}(1))(\boldsymbol{\phi}^*_{\boldsymbol{\omega}}(2))^{-1}(\boldsymbol{\phi}^*_{\boldsymbol{\omega}}(1))^T,$$

$$g(\boldsymbol{\phi}^*_{\boldsymbol{\nu}}) = -\frac{q}{2}\log(2\pi) + \frac{1}{2}\log(\det(-2(\boldsymbol{\phi}^*_{\boldsymbol{\nu}}(2)))) + \frac{1}{4}(\boldsymbol{\phi}^*_{\boldsymbol{\nu}}(1))(\boldsymbol{\phi}^*_{\boldsymbol{\nu}}(2))^{-1}(\boldsymbol{\phi}^*_{\boldsymbol{\nu}}(1))^T,$$

$$g(\boldsymbol{\phi}^*_{\upsilon^j_k}) = (\phi^*_{\upsilon^j_k}(1)+1)\log(-\phi^*_{\upsilon^j_k}(2)) - \log(\Gamma(\phi^*_{\upsilon^j_k}(1)+1)),$$

$$g(\boldsymbol{\phi}^*_{\tau_2}) = (\phi^*_{\tau_2}(1)+1)\log(-\phi^*_{\tau_2}(2)) - \log(\Gamma(\phi^*_{\tau_2}(1)+1)),$$

$$g_{a_k}(\boldsymbol{\phi}^*_{a_k}) = (\boldsymbol{\phi}^*_{a_k}(1)+1)\log(-\boldsymbol{\phi}^*_{a_k}(2)) - \log(\Gamma(\boldsymbol{\phi}^*_{a_k}(1)+1)),$$

$$g_{\alpha_i}(\boldsymbol{\phi}^*_{\alpha_i}) = (\boldsymbol{\phi}^*_{\alpha_i}(1)+1)(\log(-\boldsymbol{\phi}^*_{\alpha_i}(2))) - \log(\Gamma(\boldsymbol{\phi}^*_{\alpha_i}(1)+1)),$$

$$g(\boldsymbol{\phi}^*_{\lambda_j}) = (\phi^*_{\lambda_j}(1)+1)\log(-\phi^*_{\lambda_j}(2)) - \log(\Gamma(\phi^*_{\lambda_j}(1)+1)),$$

$$g(\boldsymbol{\phi}^*_{\lambda_2}) = (\phi^*_{\lambda_2}(1)+1)\log(-\phi^*_{\lambda_2}(2)) - \log(\Gamma(\phi^*_{\lambda_2}(1)+1)),$$

$$g(\boldsymbol{\phi}^*_{\kappa_j}) = (\phi^*_{\kappa_j}(1)+1)\log(-\phi^*_{\kappa_j}(2)) - \log(\Gamma(\phi^*_{\kappa_j}(1)+1)),$$

$$g(\boldsymbol{\phi}^*_{\chi_{2j}}) = (\phi^*_{\chi_{2j}}(1)+1)\log(-\phi^*_{\chi_{2j}}(2)) - \log(\Gamma(\phi^*_{\chi_{2j}}(1)+1)).$$

We further need the terms of the form $< \log(P[\boldsymbol{V}|\boldsymbol{H}]) >$. For the individiual level data, we have

$$L_{\boldsymbol{y}_j} = -\frac{n_j}{2}\log(2\pi) + \frac{n_j}{2}E[\boldsymbol{u}_{\lambda_j}(1)] - \frac{E[\boldsymbol{u}_{\lambda_j}(2)]}{2}(\boldsymbol{y}_j^T\boldsymbol{y}_j - 2\boldsymbol{y}_j^T\boldsymbol{X}_jE[\boldsymbol{u}_{b_j}(1)] + Tr(\boldsymbol{X}_j^T\boldsymbol{X}_j \cdot E[\boldsymbol{u}_{b_j}(2)])).$$

For the summary statistics observables we have

$$\begin{aligned} L_{\boldsymbol{z}_j} &= -\frac{m_j}{2}\log(2\pi) + \frac{m_j}{2}E[\boldsymbol{u}_{\lambda_j}(1)] - \frac{1}{2}\log(\det(\boldsymbol{\Sigma}_j)) \\ &- \frac{E[\boldsymbol{u}_{\lambda_j}(2)]}{2}(\boldsymbol{z}_j^T\boldsymbol{\Sigma}_j^{-1}\boldsymbol{z}_j - 2\sqrt{n_j}\boldsymbol{z}_j^TE[\boldsymbol{u}_{\boldsymbol{b}_j}(1)] + n_jTr(\boldsymbol{\Sigma}_jE[\boldsymbol{u}_{\boldsymbol{b}_j}(2)])). \end{aligned}$$

#### 2 Default Parameter Pettings

Default parameter settings were set to $\gamma_1 = 3$, $\tau_1 = 2$, $\phi_1 = 3$, $\phi_2 = 3$, $\xi_1 = 100$, $\xi_2 = 0.03$, $\lambda_1 = \sqrt{10^5}$, $\rho_1 = 10^5$, $\rho_2 = \sqrt{10^5}$, $p = 5$,$\chi_1 = 1$, $\chi_2 = 3 \cdot 10^{-4}$, $\zeta_1 = 1$, $\zeta_2 = 100$, $\boldsymbol{p} = \mathbf{0.2}$ and $\boldsymbol{c} = \mathbf{0}$.

### 3 Supplementary Figures

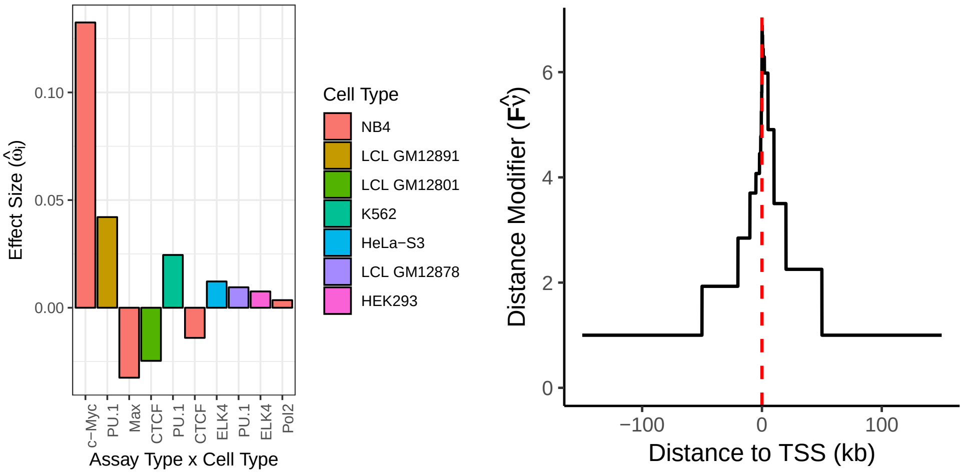


(a) (b)

Figure 1: *Parameter estimates for the directed annotation TF subset when using BAGEA on Monocyte eQTL Data.*
Shown are parameter estimates from fitting monocyte eQTL data using *TF Expecto* predictions in all cell types. a) *BAGEA* reveals the experiments underlying the directed annotations that are most predictive of gene expression. **Assay Type x Cell Type**: Each experiment is a particular assay type performed in a particular cell type. **Effect Size** ($\hat{\omega}_i$, for experiment i): The BAGEA-estimated effect on gene expression. Shown here the ten largest directed annotation effect sizes. We see *c-Myc* annotation in *NB4* dominates. b) Shown is the estimated **distance modifier** of the directed component, $\boldsymbol{F\hat{\nu}}$. We see a characteristic peak around the *TSS*, implying that the directed annotations are upweighted close to the *TSS* and more so downstream than upstream.

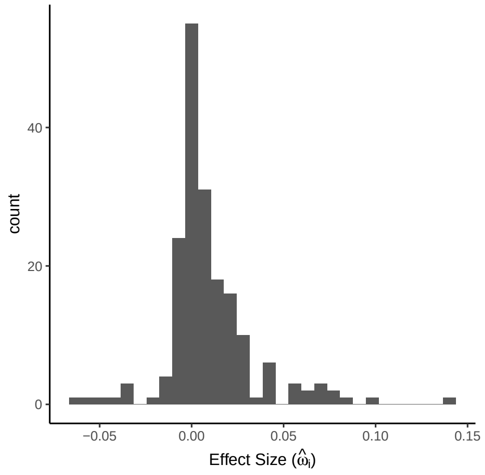


(a)

Figure 2: *Histogram of directed effect sizes $\hat{\omega}$ across all 14 GTEx data sets.*
Displayed are estimated directed annotation effect sizes $\hat{\omega}$ for all GTEx (and GEAUVADIS) data sets, with values with absolute value below $10^{-3}$ removed. Shown are results when fitting on data from all autosomes.

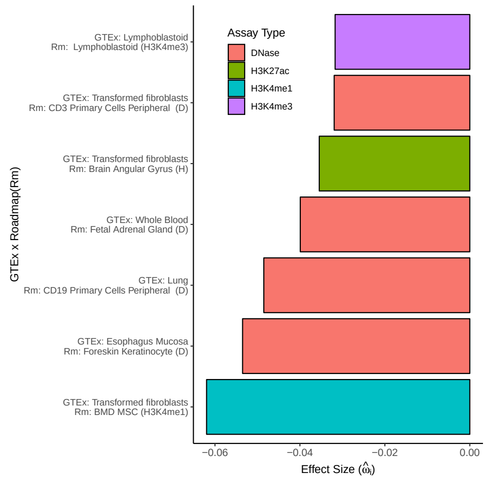


(a)

Figure 3: *Largest negative directed annotation effect sizes for GTEx summary statistics repeat some of the same tissue pairings as large positive effect sizes.*
Shown are the largest negative directed annotation effect from fitting 14 different GTEx (and GEAUVADIS) eQTL summary statistics data sets using Histone and DNase1 Expecto predictions derived from Roadmap, complementing results in Figure 6.
